# Scribble, Lgl1, and myosin IIA interact with α/β-catenin to maintain epithelial junction integrity

**DOI:** 10.1101/2023.02.15.528619

**Authors:** Maha Abedrabbo, Shirel Sloomy, Reham Abu-Leil, Einav Kfir-Cohen, Shoshana Ravid

**Affiliations:** Department of Biochemistry and Molecular Biology, The Institute of Medical Research Israel-Canada, The Hebrew University-Hadassah Medical School, Jerusalem 91120, Israel

**Keywords:** Cell Polarity, cell adhesion, EMT, tumor suppressor, adherens junction, myosin

## Abstract

E-cadherin, α- and β-catenin (E-cadherin-catenin complex) together with the cytoskeleton build the core of Adherens junctions (AJs). Scribble and Lgl1 are tumor suppressors, and it has been reported that Scribble stabilizes the coupling of E-cadherin with catenins promoting epithelial cell adhesion, but the molecular mechanism remains unknown. Here, we investigated the role of Scribble, Lgl1, and myosin-IIA (NMII-A) in AJ integrity. We show that Scribble, Lgl1, and NMII-A reside in a complex with the E-cadherin-catenin complex. Depletion of either Scribble or Lgl1 disrupts the localization of E-cadherin-catenin complex to AJs. aPKCζ phosphorylation of Lgl1 regulates AJ localization of Lgl1 and E-cadherin-catenin complex. Both Scribble and Lgl1 regulate the activation and recruitment of NMII-A at AJs. Finally, Scribble and Lgl1 are downregulated by TGFβ-induced EMT, and re-expression of Scribble or Lgl1 during EMT impedes its progression. Our results provide insight into the mechanism regulating AJ integrity by Scribble, Lgl1, and NMII-A.

## Introduction

Cell polarity lies at the heart of the establishment of proper cellular architecture and function. Epithelial cells polarize along an apical-basal axis and establish molecularly and functionally distinct apical, basal, and lateral membrane domains. At the interface between the apical and lateral domains, adherens junctions (AJs) provide mechanical strength and strong adhesive interface between cells that ensure proper tissue function ^1^. Loss of apical-basal cell polarity and AJs occurs during epithelial-mesenchymal transition (EMT) in wound healing, fibrosis, and cancer progression ^2, 3^. Epithelial polarization is controlled by an epithelial polarity program, in which polarity proteins and acto-myosin cytoskeleton, among others, play important roles ^4^. Three protein complexes establish apical-basal cell polarity: the Partitioning defective (Par) complex, the Crumbs complex, and the Scribble complex. The Scribble complex consists of Scribble (Scrib), Discs large (Dlg) and Lethal giant larvae (Lgl). In epithelial cells, the Scribble complex is located basolaterally, restricting the Par complex to the apical membrane, and *vice versa* ^5, 6^. Thus, Scribble and Par dynamically regulate apical-basal polarization by mutual exclusion. The members of the Scribble protein complex are identified as tumor suppressors ^7^. In *Drosophila*, mutations in the genes that encode these proteins caused an overgrowth of embryonic tissue, particularly in the imaginal disc and brain cells, forming large tumors ^8^. In humans, low Scrib levels associated with poor prognosis have been reported in many tumors, especially in breast and prostate cancer, confirming Scrib as a tumor suppressor ^7^. Scrib mis-localization is also associated with poor survival in prostate cancer ^9-11^, indicating the importance of Scrib localization to its functions.

Scrib is a membrane-associated scaffold protein that contains at its N-terminal a leucine-rich repeat (LRR) domain that restricts Scrib at the basolateral membrane of epithelial cells ^12^. At its C-terminal, Scrib contains 4 PDZ domains through which it interacts with many protein partners, such as β-Pix, a guanine nucleotide exchange factor (GEF) for the Rac GTPase ^13^. Scrib regulates the localization of Lgl1 at the leading edge of migrating cells and *vice versa* ^14^. There are two mammalian Lgl homologues, Lgl1 and Lgl2. Lgl1 is widely expressed, whereas Lgl2 has a more restricted expression pattern in mouse tissues ^15^. Therefore, in our studies, we focused on Lgl1. In humans, reduced expression of Lgl1 is implicated in the progression of several human tumors ^16^. Phosphorylation of Lgl at its C-terminal by the atypical protein kinase C isoform ζ (aPKCζ) regulates its cellular localization, which is important for the exclusion of the Scribble complex from the apical membrane and for the proper cell polarization of migrating cells ^6^.

Mammalian cells express three different nonmuscle-myosin-II (NMII) isoforms: NMII-A, NMII-B, and NMII-C. These isoforms play distinct but related roles in different cellular processes ^17^. NMII is a hexamer composed of two heavy chains and two pairs of essential and regulatory light chains ^17^. NMII is activated when myosin regulatory light chains (MRLC) are phosphorylated by a Ca^2+^ and calmodulin (CaM)-dependent protein kinase myosin light chain kinase (MLCK), and are inactivated when MRLC is dephosphorylated by myosin light chain phosphatase (MLCP) ^17^. Thus, MRLC phosphorylation is regulated by MLCK and MLCP, any imbalance in the coordination of the two enzymes resulting in altered migration and adhesion ^18^. NMII is essential for epithelial tissue architecture ^19-22^. It protects junctions from disassembly during development and EMT ^23^ and provides the mechanical force necessary for strengthening the AJs ^24^. In epithelial cells, NMII regulates cell-cell adhesion by determining the ability of cells to concentrate E-cadherin at contacts, and reinforce junctions and protect them from disruptive forces ^25, 26^. NMII is also required to assemble junctional complexes by activating α-catenin ^27^. Finally, during EMT, actin cytoskeleton undergoes remodeling to promote morphological changes and cell migration ^28^. In migrating cells, Lgl1 interacts directly with NMII-A, whereas Scrib seems to interact specifically with NMII-B ^14, 29^. Lgl1 regulates NMII-A filament assembly and cellular localization ^30^, and Lgl1-NMII-A interaction is regulated by phosphorylation of Lgl1 by aPKCζ ^29^.

At the core of AJs is the protein epithelial cadherin (E-cadherin) that establishes AJs through calcium-dependent homophilic binding of the extracellular cadherin domains. The cytoplasmic domain of E-cadherin directly binds to β-catenin, which in turn binds to the α-catenin that connects the AJs to the actin cytoskeleton ^31^. Scrib has been shown to interact with E-cadherin ^32^. Depletion of Scrib disrupted E-cadherin-mediated cell-cell adhesion in MDCK cells, which was partially rescued by expression of an E-cadherin-α-catenin fusion protein, indicating that Scrib stabilizes the coupling between E-cadherin and the catenins ^32^. Scrib also forms a complex with β-catenin ^33-36^, and its recruitment to E-cadherin promotes its association with p120-catenin ^32, 37^. *In vitro* and *in vivo* experiments revealed that AJ stabilization of neuroepithelium requires a direct interaction between Lgl1 and N-cadherin, and that this interaction is inhibited by aPKC-mediated phosphorylation of Lgl1 ^38^. In *Drosophila*, loss of β-catenin enhanced the phenotype observed in Lgl loss, suggesting a genetic interaction between Lgl and AJs ^39^. Thus, Scrib and Lgl1 seem to play an important role in AJs integrity. NMII-A also plays a role in regulating the localization of AJ proteins: genetic ablation of NMII-A from embryonic stem cells and mouse embryos disrupted localization of E-cadherin and β-catenin at cell-cell contacts ^40^. In MCF-7 cells, NMII-A was necessary for E-cadherin localization at cell-cell contacts, and E-cadherin was also crucial for recruitment and activation of NMII-A ^25^. NMII-A has been shown to interact with α-catenin and to be required for proper AJ assembly ^27^.

Here, we investigated the roles of Scrib, Lgl1, and NMII-A in AJ architecture. We show that depletion of either Scrib or Lgl1 disrupts the AJs localization of E-cadherin and of α-and β-catenin. In addition, we show that the localization of Lgl1 and Scrib at AJs are interdependent, as depletion of either protein affects its counterpart. We further show that Scrib and Lgl1 form a complex with the AJ proteins, E-cadherin and α- and β-catenin, through direct interaction of Lgl1 with α-catenin and of Scrib with α- and β-catenin. We also found that Lgl1 phosphorylation by aPKCζ disrupts Lgl1 localization at AJs. In addition, we provide data indicating that Scrib and Lgl1 regulate the activation and recruitment of NMII-A at AJs. Finally, we demonstrate that TGFβ−induced EMT downregulates the expression of Scrib and Lgl1, and that the re-expression of either Scrib or Lgl1 impedes TGFβ−induced EMT progression, shedding light on their roles as tumor suppressors. To our knowledge, this is the first report of downregulation of Scrib and Lgl1 expression by TGFβ-induced EMT program.

## Materials and Methods

### Cell culture and growth conditions

A549 and HEK293T cell lines were maintained in high glucose DMEM (Sigma-Aldrich) supplemented with 2 mM L-glutamine, 10% FCS and antibiotics (100 U/ml penicillin, 100 mg/ml streptomycin), and 1∶100 Biomyc3 anti-mycoplasma antibiotic solution (Biological Industries, Beit HaEmek, Israel). Cells were grown at 37°C in a humidified atmosphere of 5% CO2 and 95% air. For TGFβ stimulation, cells were cultured with 5ng/ml recombinant human TGFβ (PeproRTech) in cell medium. If cells were stimulated for more than 24 hr with TGFβ, the TGFβ in the medium was refreshed every 24 hr. For EMT induction in HMLE-Twist-ER cells, cells were treated with 10 nM 4-hydroxy tamoxifen (OHT) (Sigma-Aldrich, H7904) for the indicated number of days. During treatment, the medium was refreshed every two days.

### Antibodies

Mouse monoclonal anti-Scribble (sc-55543), anti-β-Actin (sc-69879), anti-α E-catenin (sc-9988) (for immuno-staining) and anti-E-cadherin (sc-8426) (for immunoprecipitation) were from Santa Cruz Biotechnology. Antibodies specific for the N- and C-terminal region of Lgl1 were generated in rabbits ^30^. Recombinant GFP antibodies were generated in rabbits ^41^. Antibody specific for the C-terminal region of human NMII-A was generated in rabbits according to the method of ^42^. Mouse monoclonal anti-E-cadherin (ab1416), anti-His (ab18184) and IgG (ab46540) were from Abcam. Rabbit polyclonal anti-α E-catenin (C2081) was from Sigma. Rabbit polyclonal anti-phospho-Myosin Light Chain 2 (Thr18/Ser19) (3674), anti-Lgl1 (D2B5A), anti-E-Cadherin (24E10) (for immuno-staining), anti-β-catenin (D10A8), and anti-Vimentin (D21H3) were from Cell Signaling Technology. Anti-MBP (E8032) was from New England BioLAbs.

Horseradish peroxidase-conjugated goat anti-Rabbit secondary antibody, donkey anti-Goat conjugated to Alexa Fluor 488 and goat anti-Rabbit conjugated to Cy5 were from Jackson ImmunoResearch Laboratories. Horseradish peroxidase-conjugated goat anti-Mouse (ab6789) and donkey anti-Mouse IgG H&L (Alexa Fluor 555) (ab150110) were from Abcam. All antibodies were diluted 1:1000 for Western blot and 1:100 for immunofluorescence.

### Preparation of plasmids

All restriction enzymes were from New England Biolabs. All PCR reactions were performed using Phusion High-Fidelity DNA polymerase (ThermoFisher Scientific) according to manufacturer instructions. Primers used for plasmid constructions are shown in Table 1. pGFP-Scrib was a kind gift from Patrick Humbert and Helena Richardson, La Trobe University, Melbourne, Australia. pGEX-GST-α catenin was a kind gift from Cara J. Gottardi, Northwestern University, USA. pGEX-GST-β-catenin was created with the Gibson assembly method ^43^. Briefly, pGEX-2T vector and Myc-β-catenin (a kind gift from Yinon Ben-Neriah, The Hebrew University) were subjected to PCR reaction with primers #1 and #2, #3 and #4, respectively. The PCR products were digested with *DpnI* and assembled with Gibson assembly master mix (New England Biolabs) according to the manufacturer instructions. pMBP-Lgl1 was previously described ^30^. To create p6xHis-Scrib-PDZ (701-1630aa), pGFP-Scrib was subjected to PCR with primers #5 and #6. The PCR fragment was digested with *BamHI* and *HindIII*, and the fragment was ligated into pQE-80L plasmid digested with *BamHI* and *HindIII*. To create pLenti-Neon-Lgl1-S6A and pLenti-Neon-Lgl1-S6D, we created phospho-mutants pmCherry-Lgl1-S6A and S6D, in which all six serine residues 650, 654, 658, 662, 669, and 672 were mutated to alanine (Ala) or aspartate (Asp), respectively, in two steps. First, we created pmCherry-Lgl1-S4A and S4D, where serine residue 650 was changed to alanine or aspartate respectively, in addition to residues 654, 658, and 662 that were changed in previously described pGFP-Lgl1-S3A and S3D ^29^. pGFP-Lgl1-S3A and S3D were subjected to two PCR reactions, first PCR with primer #7 and #8, #7 and #9, respectively; second PCR with primers #10 and #11, #10 and #12, respectively. PCR products were used as a template for a third PCR with primers #7 and #9. The third PCR products were digested with *BamHI* and *EcoRI*, and the fragments were ligated into pmCherry plasmid digested with *EcoRI* and <u>BamHI</u>. The final plasmid was pmCherry-Lgl1-S4A and pmCherry-Lgl1-S4D. To change all six mentioned serines, we create pmCherry-Lgl1-S6A and pmCherry-Lgl1-S6D, pmCherry-Lgl1-S4A and pmCherry-Lgl1-S4D were subjected to two PCR reactions, the first with primer #7 and #11, #7 and #12, respectively, and the second with primers #10 and #15, #10 and #16, respectively. The PCR products were inserted into pmCherry plasmid as described above. To create our final plasmid pLenti-Neon-Lgl1-S6A and pLenti-Neon-Lgl1-S6D, pmCherry-Lgl1-S6A and pmCherry-Lgl1-S6D, described above, were subjected to PCR reaction with primers #17 and #10. PCR fragments were digested with *BamHI* and ligated into pLenti-Neon plasmid digested with *BamHI*.

**Table 1:**
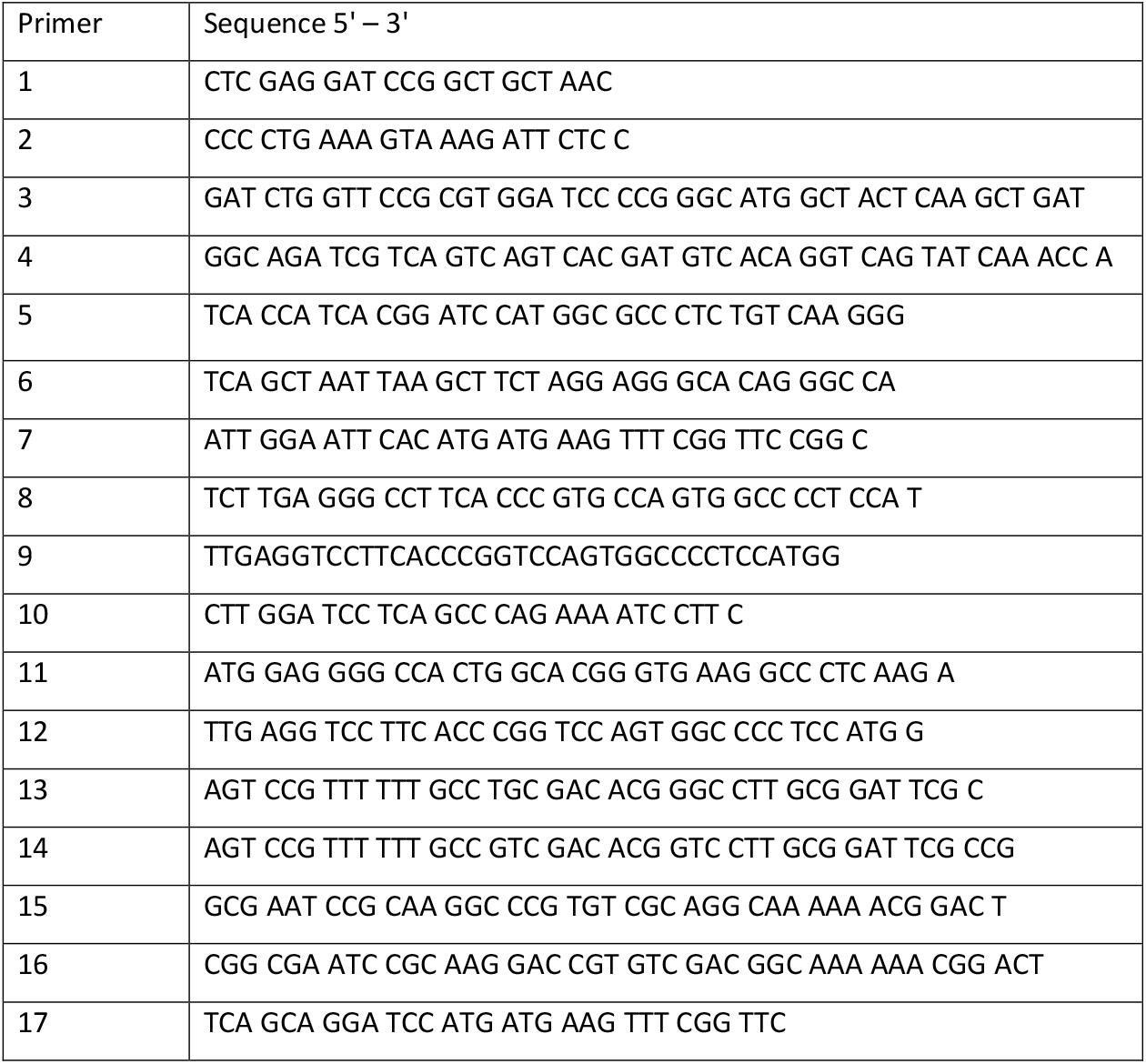
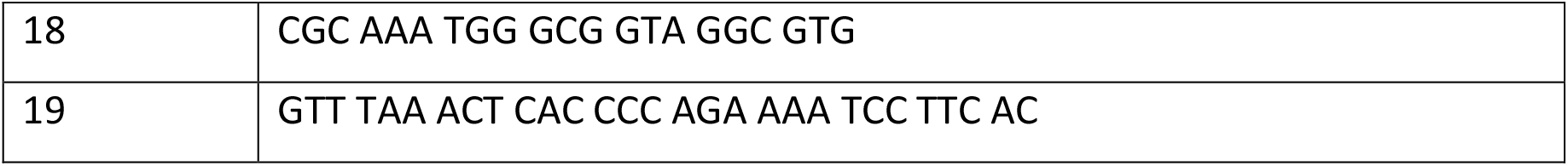
primers used in this study

pLKO-Tet-shscrib, pLKO-Tet-shLgl1 and pLenti-Neon-Lgl1-WT were previously described ^14^. To create the inducible GFP–Scrib expression vector (pVPR-GFP-Scrib), pGFP–Scrib and PB-TRE-dCas9-VPR (Addgene plasmid # 63800) ^44^ were digested with *BamHI* and *PmeI*, respectively, followed by fill-in reaction using Klenow fragments and digestion with *BmtI*. pGFP-Scrib fragments were ligated into PB-TRE-dCas9-VPR. To create an inducible GFP–Lgl1 expression vector (pVPR-GFP-Lgl1), pGFP-Lgl1 ^30^ was subjected to PCR with primers #18 and #19. PCR products were digested with *BmtI* and *PmeI* and ligated into PB-TRE-dCas9-VPR digested with *BmtI* and *PmeI*.

### Generation of inducible Scrib and Lgl1 depleted cell lines

HEK293T cells grown to 90% confluency were transfected with pRev, pPM2, pTat, pVSVG, and pLKO-Tet-shScrib (target 3UTR) or pLKO-Tet-shLgl1 (target 3UTR) using PEI; 8 h posttransfection, the medium was replaced with fresh medium. At 24 and 48 hr posttransfection, the supernatant was collected and filtered with a 0.45-μm filter, and 4 μg/ml polybrene was added. A549 cells were infected with viruses containing pLKO-Tet-shScrib or pLKO-Tet-shLgl1; 24 hr post-infection, the medium was replaced with fresh medium, and 48 hr post-infection, fresh medium containing 2 µg/ml puromycin (Sigma-Aldrich) was added to the infected and control cells (uninfected A549 cells). After the control cells died, knockdown of Scrib or Lgl1 was induced with 1 µg/ml doxycycline (Dox); after Dox addition, Scrib and Lgl1 knockdown was detected by Western blot after 72 and 24 hr, respectively (Figure S1).

### Generation of inducible GFP-Scrib (Scrib-Rescue) and GFP-Lgl1 expressing cell lines

HEK293T cells grown to 90% confluency were transfected with pRev, pPM2, pTat, pVSVG, and pVPR-GFP-Scrib or pVPR-GFP-Lgl1 using PEI; 8 hr posttransfection, the medium was replaced with fresh medium. At 24 and 48 hr posttransfection, the supernatant was collected and filtered with a 0.45-μm filter, and 4 μg/ml polybrene was added. A549-tet-shScrib and A549 cells were infected with viruses containing inducible GFP-Scribble or GFP-Lgl1, respectively; 24 hr post infection, the medium was replaced with fresh medium. GFP expression was induced with 1 µg/ml doxycycline (Dox); after Dox addition, GFP positive cells were isolated by sorting using BD FACSAria III. A549-tet-shScrib-VPR-GFP-Scrib cells were treated with Dox before rescue experiments, ensuing knock-down of endogenous Scrib.

### Generation of Neon-Lgl1-WT (Lgl1-Rescue) and phosphomutant Lgl1 cell lines

HEK293T cells grown to 90% confluency were transfected with pRev, pPM2, pTat, pVSVG, and pLenti-Neon-Lgl1-WT, pLenti-Neon-Lgl1-S6A or pLenti-Neon-Lgl1-S6D using PEI; 8hr posttransfection, the medium was replaced with fresh medium. At 24 and 48 hr posttransfection, the supernatant was collected and filtered with a 0.45-μm filter, and 4 μg/ml polybrene was added. A549-tet-shLgl1 cells were infected with viruses containing pLenti-Neon-Lgl1-WT, pLenti-Neon-Lgl1-S6A or pLenti-Neon-Lgl1-S6D; 24 hr post-infection, the medium was replaced with fresh medium and neon positive cells were isolated by sorting using BD FACSAria III. Cells were treated with Dox before all experiments, ensuing knock-down of endogenous Lgl1.

### Co-immunoprecipitation assays

A549 cells were grown in a 100-mm dish to 90% confluency. After 18 hr, A549 cells were used for co-IP assay of endogenous proteins. For A549 expressing GFP-only, A549 cells were transfected with 12 µg of pEGFP-C2 mixed with 36 µg PEI in DMEM and harvested after 24 hr hours. Cells were harvested with 800µl NP-40 buffer (20mM Tris pH=8, 150-200mM NaCl, 0.5mM EDTA, 1% NP-40, 1mM DTT, 5% glycerol and protease inhibitor cocktail; Sigma-Aldrich). Protease inhibitor cocktail (Bimake, #B15002) was added to buffer in experiments precipitating with anti E-cadherin. Cell extracts were sonicated and centrifuged at 4°C for 15 min at 16,000 ×g. The appropriate antibodies were incubated with cell lysate on a rotator at 4°C for 2 h. The lysate-antibodies mix was added to A/G beads prewashed with 300μl NP-40 buffer and incubated on a rotator at 4°C for 2 h. Then, the mix was washed three times with NP-40 buffer and SDS-sample buffer was added. Samples were dissolved on SDS-PAGE and analyzed by Western blot. Proteins from Western blots were detected using the EZ-ECL Chemiluminescence Detection kit (Biological Industries), and the band intensity was analyzed using Bio-Rad Gel dox CR Luminescent Image Analyzer and Fujifilm Image Gauge Ver. 3.46 software (Fujifilm, Tokyo, Japan). The ImageGauge software detects saturation to ensure the linearity of the signal.

### Protein expression and purification

p6xHis-Scrib-PDZ was grown in T7+ strain *E. coli* at 37°C to an optical density at 600 nm (OD600nm) of 0.5, then 0.5 mM IPTG was added and the bacteria were grown at 16°C overnight. Bacterial pellets were collected by centrifugation at 4°C for 15 min at 10,000 xg (Sorvall, Thermo Scientific, Rotor F12) and resuspended in buffer A (50mM Tris pH=8, 500mM NaCl, 20mM imidazole, 20mM β-mercaptoethanol) and 0.5 mM phenylmethylsulfonyl fluoride (PMSF) and 1% Tween-20. Bacteria suspensions were sonicated and centrifuged using F21-8×50y rotor (Thermo-Scientific) at 4°C for 15 min at 20,000 xg. Supernatants were collected and loaded on a Ni^2+^-NTA bead column (GE Healthcare) prewashed with buffer A. His-Scrib-PDZ protein was eluted with buffer A containing 250 mM imidazole. Fractions containing protein were pooled and dialyzed against buffer A without imidazole and 150mM NaCl. Protein concentration was determined by comparing the densitometry of the band of a protein sample on Coomassie-stained SDS-PAGE gel to known amounts of bovine serum albumin (BSA) samples on the same gel. MBP-Lgl1 was purified as described previously ^30^. MBP-Lgl1 was eluted with 10mM maltose with MBP buffer.

pGEX-2T, pGEX-GST-α-catenin and pGEX-GST-β-catenin were grown in T7+ strain *E. coli* at 37°C to an optical density at 600 nm (OD600nm) of 0.5, then 0.5 mM IPTG was added and the bacteria were grown at 16°C overnight. Bacterial pellets were collected by centrifugation at 4°C for 15 min at 10,000 g (Sorvall, Thermo Scientific, Rotor F12) and resuspended in lysis buffer (50mM Tris pH=8, 150mM NaCl, 5mM EDTA, 1mM DTT, 1% TX-100 and 1mM fluoride PMSF) and 1mg/ml lysozyme, and incubated on ice for 15 min. Bacterial suspensions were sonicated and centrifuged in a F21-8×50y rotor (Thermo Scientific) at 4°C for 15 min at 20,000 xg. Supernatants were collected and mixed with 500µl glutathione beads (Rimon Biotech) prewashed with lysis buffer without PMSF. The bacterial lysates and glutathione beads were incubated at 4°C on a rotator for 2hr followed by three washes with lysis buffer. GST protein concentration was determined as described above.

### Pull-down assays

GST-only, GST-α-catenin or GST-β-catenin proteins immobilized on glutathione beads were washed three times with binding buffer (20mM Tris pH=8, 200mM NaCl, 10% glycerol, 1% TX-100, 1mM DTT), then incubated with 10µg MBP-Lgl1 or His-Scrib-PDZ in a final volume of 200µl for 1hr on rotator at 4°C. Beads were washed three times with binding buffer and eluted with 30µl elution buffer (50mM Tris pH=9, 150mM NaCl, 20mM Glutathione, 30mM imidazole) for 1 h on rotator at 4°C. The beads were centrifuged at 4°C for 5 min at 16000 xg, and 25µl of the supernatant were added to 25µl SDS sample buffer. For total protein input, 15µl of the beads-proteins mix were added to 15µl of SDS sample buffer. Proteins were analyzed on SDS-PAGE gels and detected with Ponceau S staining solution or Western blot.

### Immunofluorescence

Cells were seeded on coverslips coated with Collagen type I (Sigma-Aldrich). After 14 h, cells were fixed with 4% paraformaldehyde in PBS for 15 min, washed three times with PBS, and permeabilized for 3 min with PBS containing 0.2% TX-100 and 0.5% BSA. Cells were washed three times and blocked with horse serum diluted 1:50 for 35 min at 37°C. Cells were washed and incubated with primary antibodies in PBS containing 0.1% BSA overnight at 4°C. Coverslips were washed three times and incubated with secondary antibodies in PBS containing 0.1% BSA for 1hr at room temperature. Coverslips were mounted on slides (Thermo-Scientific) using Vectashield mounting medium (Vector Laboratories). Confocal images were obtained with Nikon Confocal A1R. Signal intensity of Scrib, Lgl1, E-cadherin, α- and β-catenin were measured with the ImageJ software package (National Institutes of Health, Bethesda, MD). For co-localization analysis, the PCC was calculated between the intensity profiles of Scrib, Lgl1, and E-cadherin at cell-cell contact. For Junctional/Cytoplasmic intensity analysis, signal intensity was measured at AJ (junctional) and cytoplasm (cytoplasmic), and average measurements at each point of junctional intensity were divided by cytoplasmic intensity for indicated proteins. The PCC and signal intensity were calculated using Excel (Microsoft) and Prism 6 (GraphPad). Statistical analysis was performed using Prism 6. Data were examined by ANOVA with a *post hoc* test between the groups. For Z-stack imaging, optical sections were collected at 400 nm intervals. Image analysis and reconstitution were carried out using NIS-Elements AR.

For pMRLC staining, cells were stimulated with 5ng/ml TGFβ for 30 min before fixation, and slides were processed as described above. For TGFβ-induced EMT staining, cells were stimulated with 5ng/ml TGFβ for 16 hr before fixation, and slides were processed as described above. For re-expression of Scrib and Lgl1 to revert EMT staining, cells were stimulated with 5ng/ml TGFβ for 24 hr, then 1 µg/ml Dox was added for additional 48 hr with or without TGFβ.

### Phosphorylation levels of MRLC

5×10^5^ cells were seeded on 30-mm dishes. After 16 hr, cells were stimulated with 5ng/ml TGFβ for 30 min, washed and SDS sample buffer was added. All samples were incubated for 5 min at 90°C, separated on 15% SDS-PAGE and analyzed with Western blot. Band intensity was analyzed as described above. Relative phosphorylation levels were quantified by dividing the intensity of the phosphorylated MRLC band by the intensity of the Actin band using Excel (Microsoft). Data were examined by two-tailed Student’s *t* test between each group.

### Protein expression following EMT induction

A549 or HMLE-Twist-ER cells were induced with TGFβ or OHT, respectively for the indicated point times and lysed with 100μl extraction buffer (50mM Tris pH=8.0, 150 mM NaCl, 0.5mM EDTA, 1% NP-40, and 1mM DTT, and protease inhibitor cocktail [Sigma-Aldrich]). Cell extracts were sonicated and centrifuged at 4°C for 15 min at 16,000 ×g. SDS sample buffer was added to supernatant. All samples were incubated for 5 min at 90°C, separated on 8% SDS-PAGE and analyzed with Western blot. Band intensity was analyzed as described above. Relative protein expression was quantified by dividing the intensity of the indicated protein band by the intensity of the actin band, using ImageGauge software and Excel (Microsoft) as described above. Data were examined by two-tailed Student’s *t* test between each group.

### Re-expression of Scrib and Lgl1 to revert EMT

1×10^6^ cells were seeded on 60-mm dishes and stimulated with 5ng/ml TGFβ for 24 hr, then 1 µg/ml Dox was added for additional 48 hr with or without TGFβ. Cells were harvested with 300μl extraction buffer (50mM Tris pH=8, 150mM NaCl, 0.5mM EDTA, 1% NP-40, 5%, 1mM DTT, and protease inhibitor cocktail [Sigma-Aldrich]). Cell extracts were sonicated and centrifuged at 4°C for 15 min at 16,000 ×g. All samples were incubated for 5 min at 90°C, separated on 8% SDS-PAGE and analyzed with Western blot. Band intensity was analyzed as described above. To calculate the extent of E-cadherin expression, the intensity of E-cadherin with TGFβ was divided by the intensity of E-cadherin without TGFβ relative to Actin, using Excel (Microsoft). Data were examined by two-tailed Student’s *t* test between each group.

For phase-contrast images, 5×10^5^ cells were seeded on 30-mm dishes and stimulated with 5ng/ml TGFβ for 24 hr, then 1 µg/ml Dox was added for an additional 48 hr with or without TGFβ. Phase-contrast of randomly chosen fields were taken with Nikon Eclipse Ts2 at ×20.

## Results

### Lgl1 and Scrib co-localize with E-cadherin at AJs

In order to begin to unravel the role of Lgl1 and Scrib in AJ integrity, we first characterized the cellular localization of Lgl1, Scrib, and E-cadherin in monolayers of A549 cells a lung carcinoma cell line, which has an epithelial origin and expresses the proteins that are the focus of this study. For this purpose, we immunostained confluent monolayers of A549 cells for E-cadherin, Lgl1 and Scrib and acquired z-stack images. Both endogenous Scrib and Lgl1 co-localized with E-cadherin at AJs (Figure 1A-C). To analyze the level of Scrib, Lgl1, and E-cadherin co-localization, we calculated the Pearson correlation coefficient (PCC) between the fluorescence intensity profiles of Scrib, Lgl1, and E-cadherin at AJs. Quantitatively, the PCC between Scrib and E-cadherin and between Lgl1 and E-cadherin was 0.83 ± 0.1 and 0.9 ± 0.08, respectively (Figure 1B). These results further verify that Scrib and Lgl1 co-localized with E-cadherin. The co-localization of Scrib and E-cadherin is consistent with previously published results ^35, 37, 45^.

**Figure 1:**
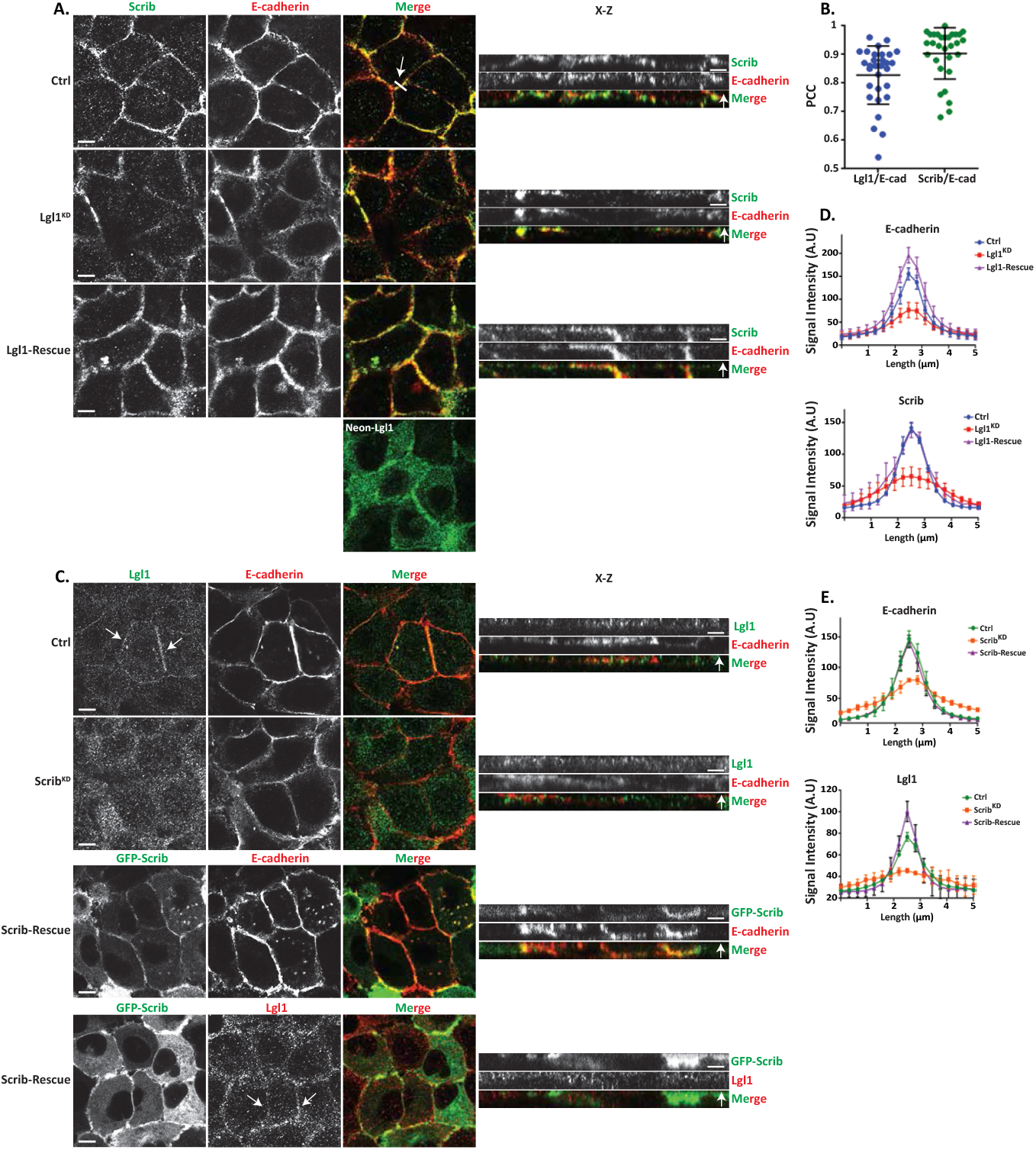
Lgl1 and Scrib regulate E-cadherin localization at AJs. **(A)** A549 tet-shLgl1 cells without (Ctrl) and with Dox (Lgl1^KD^) and A549 tet-shLgl1-Neon-Lgl1 with Dox (Lgl1-Rescue) were fixed and immunostained for Scrib and E-cadherin. Neon-Lgl1 only is shown at the bottom of A. Left, representative images from Z-stack of E-cadherin and Scrib from the apical side, scale bar, 10 µm. White arrow points at the line used to measure the signal of fluorescence intensity at AJs presented in D. Right, Z-stack constructed from serial optical sections along apico-basal axis. White arrow indicates apical direction along apico-basal axis in Z-stack. Scale bar, 5 µm. **(B)** Pearson correlation coefficient (PCC) between the fluorescence intensity of endogenous E-cadherin and Scrib or Lgl1. Results are mean ± SD, n = 30. **(C)** A549 tet-shscrib without (Ctrl) and with Dox (Scrib^KD^) and A549 tet-shScrib-GFP-Scrib with Dox (Scrib-Rescue) cell lines were fixed and immunostained for Lgl1 and E-cadherin. Left, representative images from Z-stack of E-cadherin and Lgl1 from the apical side, scale bar, 10 µm. Arrows indicate Lgl1 localization at AJs. Right, Z-stack obtained as described above. White arrow indicates apical direction along apico-basal axis in Z-stack. Scale bar, 5 µm. (**D)** Fluorescence intensity of endogenous E-cadherin or Scrib measured at AJs in the indicated cell lines. Results are mean ± SD, for E-cadherin n= 45-80, for Scrib n=60. **(E)** Fluorescence intensity of endogenous E-cadherin or Lgl1 measured at AJs in indicated cell lines. Results are mean ± SD, for E-cadherin n=50-70, for Lgl1 n=60. A.U.: arbitrary units.

### Depletion of Lgl1 or Scrib disrupts the integrity of AJ proteins

To assess the functional role of Lgl1 and Scrib in AJ architecture, we generated A549 depleted of either Lgl1 (Lgl1^KD^) or Scrib (Scrib^KD^). Lgl1 and Scrib protein levels were significantly reduced in Lgl1^KD^ and Scrib^KD^ cells, respectively (Figure S1A-B). Depletion of Lgl1 did not affect the expression levels of Scrib, and *vice versa* (Figure S1A-B). Furthermore, depletion of either Lgl1 or Scrib did not affect the expression levels of E-cadherin-catenin proteins (i.e., E-cadherin, α-catenin, and β-catenin) (Figure S1A-B). To test the effect of Lgl1 or Scrib depletion on AJ architecture, we analyzed the morphology of AJs in Lgl1^KD^ and Scrib^KD^ cells by immunofluorescence microscopy and acquired z-stack images. We found that depleting either Lgl1 or Scrib expression led to a significant loss of E-cadherin from the AJs, and E-cadherin exhibited a diffusion pattern at the cell cortex (Figure 1A-C). The loss of E-cadherin at junctions was not due to a decrease in E-cadherin protein levels (Figure S1). Note that E-cadherin decrease at AJs was not uniform in Lgl1^KD^ and Scrib^KD^ monolayer cells, probably reflecting the differences in knockdown efficiency in the stable mixed population of Lgl1^KD^ and Scrib^KD^ cells. Quantification of E-cadherin fluorescence intensity across the AJs in Lgl1^KD^ and Scrib^KD^ cells showed a large decrease in E-cadherin intensity peak at the junctions (Figures 1D-E, S2 and S3). Together, these results indicate that AJ integrity is compromised when Lgl1 or Scrib expression is reduced. The results of AJ disruption in the absence of Scrib are consistent with previous results using MDCK cells ^32^. Thus, Lgl1 and Scrib are required for the proper junctional localization of E-cadherin.

We have previously shown that the cellular localizations of Lgl1 and Scrib in migrating cells is interdependent ^14^. To test whether the junctional localizations of Lgl1 and Scrib is also interdependent, we stained Scrib and Lgl1 in Lgl1^KD^ and Scrib^KD^ cell lines, respectively. We found that Scrib in Lgl1^KD^ cells and Lgl1 in Scrib^KD^ cells were significantly lost from AJs (Figure 1A-C). These results were further supported by quantification of fluorescence intensity across the AJs of Scrib and Lgl1 in Scrib^KD^ and Lgl1^KD^ cell lines, respectively (Figures 1D-E, S2 and S3). Thus, similarly to migrating cells, Lgl1 and Scrib junctional localization is interdependent. To further analyze the effect of Lgl1 and Scrib depletion on AJ architecture, we examined the junctional localization of α- and β-catenin in Lgl1^KD^ and Scrib^KD^ cell lines. We found that similarly to E-cadherin, α- and β-catenin were lost from AJs (Figure 2A-B), and quantification of fluorescence intensity across the AJs in Lgl1^KD^ and Scrib^KD^ cells showed a large decrease in α- and β-catenin intensity peak (Figure 2C-D, S2 and S3).

**Figure 2:**
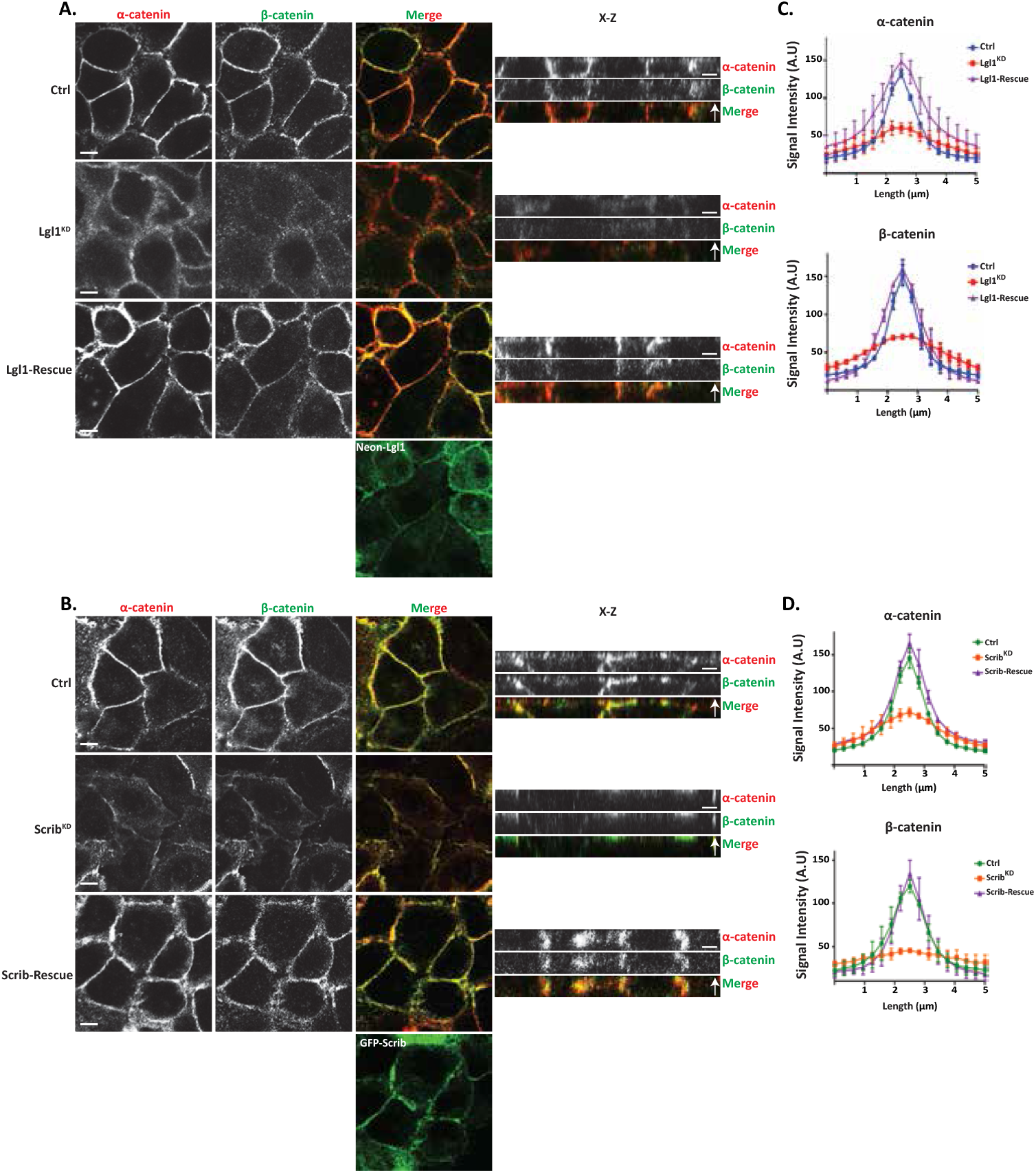
Lgl1 and Scrib regulate α- and β-catenin localization at AJs. **(A)** and **(B)** A549 tet-shLgl1 without (Ctrl) and with Dox (Lgl1^KD^) and A549 tet-shLgl1-Neon-Lgl1 with Dox (Lgl1-Rescue), A549 tet-shscrib without (Ctrl) and with Dox (Scrib^KD^), and A549 tet-shScrib-GFP-Scrib with Dox (Scrib-Rescue) cell lines were fixed and immunostained for α- and β-catenin. Left, representative images from Z-stack of α- and β-catenin from the apical side, scale bar, 10 µm. Right, Z-stack constructed from serial optical sections along apico-basal axis. Scale bar, 5 µm. **(C)** and **(D)** Fluorescence intensity of endogenous α- and β-catenin measured at AJs in the indicated cell lines as described in Figure 1. Results are mean ± SD, for n=60 for α- and β-catenin in all Lgl1 cell lines. In Scrib cell lines, results are mean ± SD, for α-catenin n=60 and for β-catenin n=35-60. A.U.: arbitrary units.

Re-expression of Lgl1 and Scrib in Lgl1^KD^ and Scrib^KD^ cells, respectively, restored the junctional localization of E-cadherin and of α- and β-catenin (Figures 1, 2, S2 and S3), demonstrating that the aberrant cellular localization of these proteins was specifically caused by the loss of Lgl1 or Scrib. Thus, Lgl1 and Scrib are important for the junctional localization of E-cadherin and of α- and β-catenin, and for maintaining AJ integrity. Note that re-expression of either mNeon-Lgl1 and GFP-Scrib did not affect the expression levels of E-cadherin-catenin complex proteins (Figure S1C-D). Similarly, re-expression of Lgl1 in Lgl1^KD^ cells and of Scrib in Scrib^KD^ cells restored the AJ localization of Scrib and Lgl1, respectively (Figures 1, 2, S2 and S3).

### Lgl1 and Scrib form a complex with AJ proteins through direct interactions with α- and β-catenin

To further explore the role of Lgl1 and Scrib in AJ integrity, we characterized the complex formation of Lgl1 and Scrib with E-cadherin-catenin proteins in A549 cells. It was previously shown that Scrib forms a complex with β-catenin ^33-36^, and Lgl1 forms a complex with N-cadherin ^38^, a highly similar protein to E-cadherin ^46^. Together, suggesting that Scrib and Lgl1 form complexes with the E-cadherin-catenin proteins to stabilize AJs. To test this hypothesis, we performed an array of immunoprecipitation assays using monolayers of A549 cells. We found that both endogenous and exogenous Scrib co-immunoprecipitated with endogenous Lgl1 and E-cadherin-catenin proteins (Figures 3A and S4A) and endogenous Lgl1 co-immunoprecipitated with endogenous α-catenin and Scrib (Figure 3B). In reciprocal experiments, we found that endogenous E-cadherin co-immunoprecipitated with exogenous Scrib, as well as α- and β-catenin (Figure 3C) and endogenous α-catenin co-immunoprecipitated with endogenous Lgl1 and with E-cadherin and β-catenin (Figure 3D). Thus, Scrib and Lgl1 form complexes with E-cadherin-catenin proteins. Next, we sought to determine whether Scrib interacts directly with α- and β-catenin. It has been shown that Scrib forms a complex with β-catenin through its PDZ domain ^47^, to test this we used His-tagged PDZ domain of Scrib (Scrib-PDZ) and GST-β-catenin recombinant proteins. As shown in Figure 3E, the Scrib-PDZ domain interacts directly with β-catenin. Next, we examined whether Scrib-PDZ also interacts directly with α-catenin. As shown in Figure 3F, Scrib-PDZ interacts with GST-α-catenin, therefore, the Scrib-PDZ domain interacts with both α- and β-catenin. Next, we tested whether Lgl1 interacts directly with α- and β-catenin. To this end, we performed a pull-down assay using recombinant Lgl1 fused to maltose-binding protein (MBP) and GST-α- or -β-catenin. We found that MPB-Lgl1 but not MBP only binds specifically to GST-α-catenin (Figure 3G), indicating that Lgl1 interacts directly with α-catenin. However, we could not detect a direct interaction between MBP-Lgl1 and GST-β-catenin (Figure S4B). Together, these results indicate that Scrib and Lgl1 form a complex with cadherin-catenin proteins through direct interactions of Scrib with α-and β-catenin, and of Lgl1 with α-catenin (Figure 3H). To our knowledge, this is the first study presenting a multi-protein complex of Scrib, Lgl1 and cadherin-catenin proteins.

**Figure 3:**
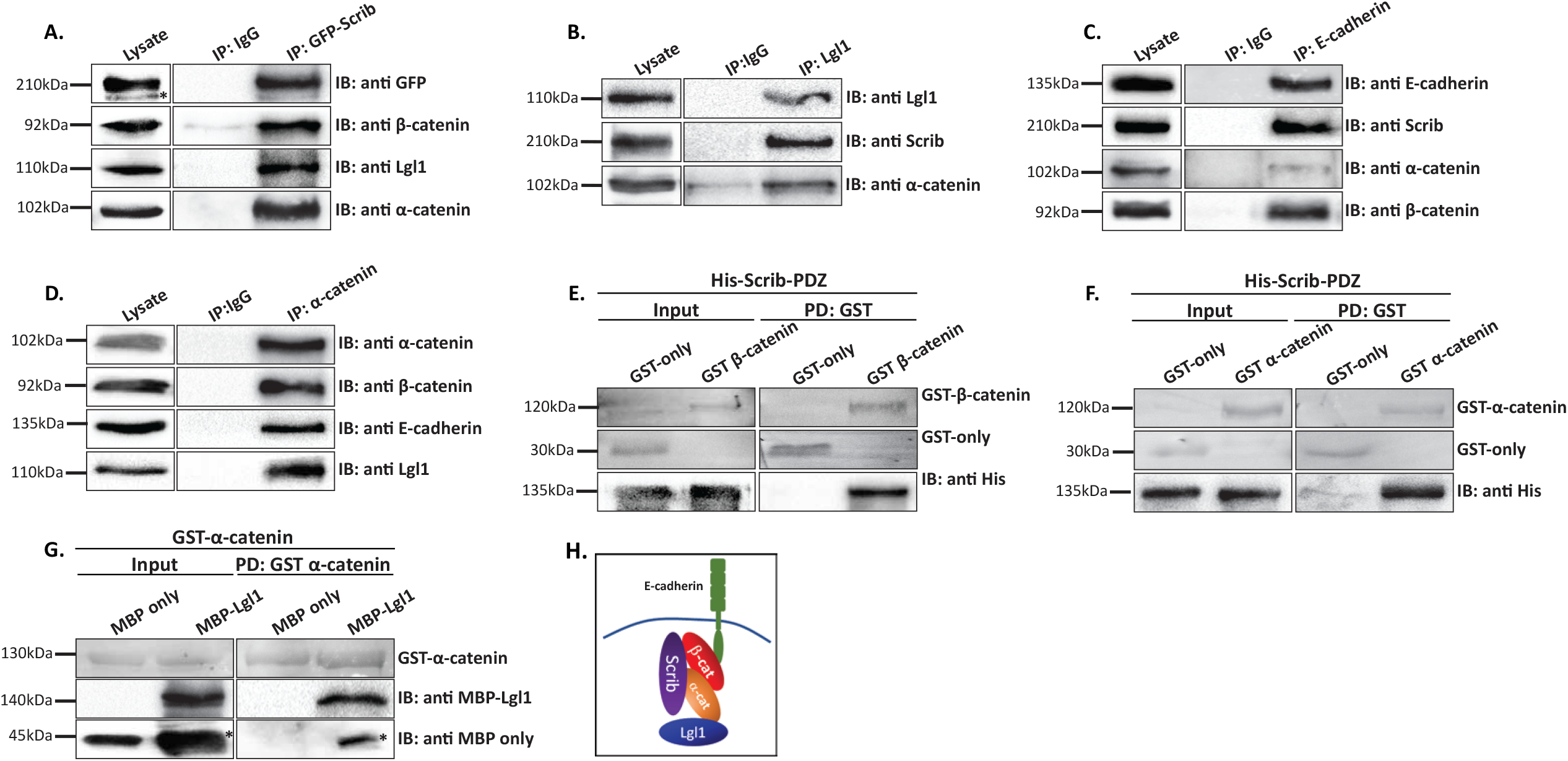
Lgl1, Scrib, and NMII-A interact with E-cadherin-catenin proteins. **(A)** A549 cells stably expressing GFP-Scrib were subjected to co-immunoprecipitation (co-IP) assay using GFP antibody. The immunoprecipitated proteins were analyzed by immunoblotting (IB) with antibodies against GFP, Lgl1, α- and β-catenin. IgG antibody was used as a negative control. **(B)** A549 cell extracts were subjected to co-IP assay using Lgl1 antibody. The immunoprecipitated proteins were analyzed by IB with antibodies against Scrib, Lgl1 and α-catenin. IgG antibody was used as negative control. **(C)** A549 cells stably expressing GFP-Scrib were subjected co-IP assay using E-cadherin antibody. The immunoprecipitated proteins were analyzed by IB with antibodies against E-cadherin, Scrib, α- and β-catenin. IgG antibody was used as a negative control. **(D)** A549 cell extracts were subjected to co-IP assay using α-catenin antibody. The immunoprecipitated proteins were analyzed by IB with antibodies against E-cadherin, Lgl1, α- and β-catenin. IgG antibody was used as negative control. **(E)** and **(F)** His-Scrib-PDZ, GST only, and GST-α-catenin (E) or GST-β-catenin (F) were subjected to pull-down (PD) assay. His-Scrib-PDZ was analyzed by IB with antibody against His-tag, and GST proteins were analyzed by Ponceau S staining. **(G)** GST-α-catenin and MBP only or MBP-Lgl1 were subjected to PD assay. MBP proteins were analyzed by IB with antibody against MBP-tag, and GST-α-catenin was analyzed by Ponceau S staining. **(H)** A model depicting the protein complex formed by Lgl1, Scrib, E-cadherin, α- and β-catenin at AJs. *Non-specific band. Molecular weights of the proteins are indicated.

### Lgl1 phosphorylation plays a regulatory role in maintaining the integrity of AJ proteins

aPKCζ phosphorylates Lgl, regulating its spatial localization in *Drosophila* epithelial cells and in migrating fibroblast cells ^29, 48, 49^. Thus, we hypothesized that Lgl1 phosphorylation by aPKCζ also affects its localization at AJs. To test this hypothesis, we created phospho-Lgl1 mutants by replacing the six serine residues phosphorylated by aPKCζ, i.e., 641, 645, 649, 653, 660, and 663 ^48^ with either alanine residues to create phospho-resistant Lgl1 (Lgl1-S6A), or with aspartate residues to create phosphomimetic Lgl1 (Lgl1-S6D). Next, we created A549 cell lines that are depleted of Lgl1 and stably expressing either Neon-Lgl1-WT, Neon-Lgl1-S6A, or Neon-Lgl1-S6D (Figure S1C). To study the effect of Lgl1 phosphorylation on its AJ localization and the localization of Scrib and E-cadherin-catenin proteins, we immunostained confluent monolayers of a phosphomimetic Lgl1 cell line for these proteins and acquired z-stack images. We found that unlike Neon-Lgl1-WT, Neon-Lgl1-S6D was diffused throughout the cell, as well as Scrib, E-cadherin, α- and β-catenin localization at AJs was disrupted (Figure 4A-B). Quantification of the fluorescence intensity of these proteins across AJs further indicates that expression of Neon-Lgl1-S6D was unable to rescue the cellular localization of E-cadherin-catenin proteins or of Scrib in comparison to Neon-Lgl1-WT (Figure 4C-D). Thus, Lgl1 phosphorylation leads to a localization pattern of adherens junction proteins that is similar to that of cells depleted of Lgl1, mimicking the phenotype of cells depleted of Lgl1, verifying the significance of Lgl1 localization at AJ and the notion that Lgl1’s function stems from its localization.

**Figure 4:**
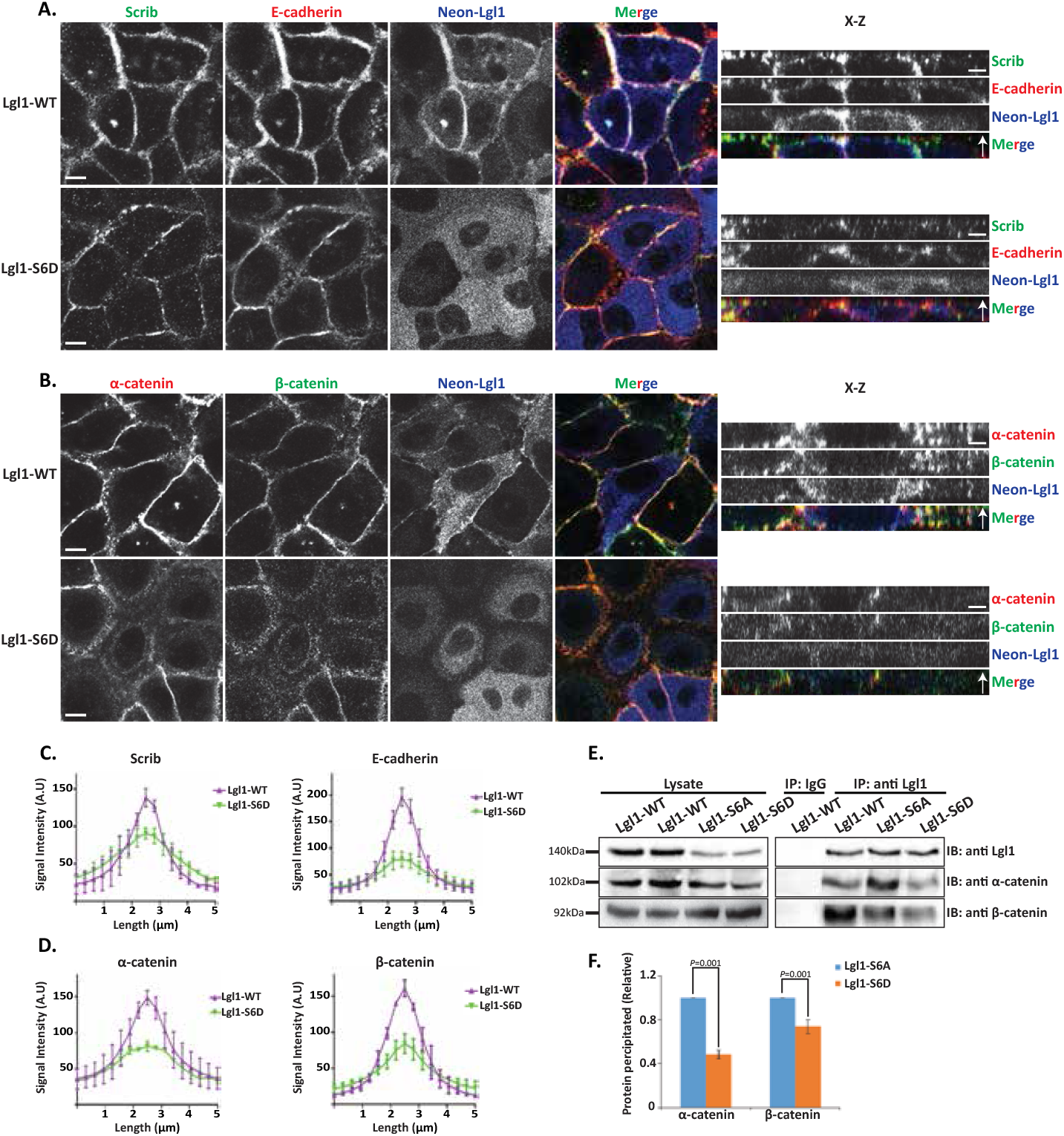
Lgl1 phosphorylation regulates AJ integrity. **(A)** and **(B)** A549-tet-shLgl1 (with Dox) cells expressing Neon-Lgl1-WT (Lgl1-WT) or Neon-Lgl1-S6D (Lgl1-S6D) were fixed and immunostained for Scrib and E-cadherin (A) or α- and β-catenin (B). For clarity, Neon-Lgl1 proteins are shown in blue. Left, representative images from Z-stack of indicated proteins from the apical side, scale bar, 10 µm. Right, Z-stack constructed from serial optical sections along apico-basal axis. Scale bar, 5 µm. **(C)** and **(D)** Fluorescence intensity of endogenous Scrib and E-cadherin, α- and β-catenin measured at AJs in the indicated cell lines as described in Figure 1. Results are mean ± SD, for Scrib n=60, for E-cadherin n= 45-80, for α- and β-catenin n=60. A.U.: arbitrary units. **(E)** Extracts from A549 cells expressing Neon-Lgl1-WT, Neon-Lgl1-S6A, or Neon-Lgl1-S6D were subjected to immunoprecipitation assay using Lgl1 antibody. The immunoprecipitated proteins were analyzed by IB with antibodies against Lgl1 and α- and β-catenin. IgG antibody was used as negative control. **(F)** Quantification of the relative amounts of α- and β-catenin precipitated by Neon-Lgl1-S6A or Neon-Lgl1-S6D. Since Lgl1-WT includes both the phosphorylated and un-phosphorylated Lgl1, the quantification of the extent of Lgl1-S6D immunoprecipitated with α- and β-catenin is relative to that of Lgl1-S6A. Values are the mean ± SD from three independent experiments subjected to two-tailed, two-sample, and unequal-variance Student’s *t* test. Molecular weights of the proteins are indicated.

The finding that expression of Neon-Lgl1-S6D disrupts the AJ localization of α- and β-catenin suggests that phosphorylation of Lgl1 regulates its complex formation with these proteins. To explore this hypothesis, we immunoprecipitated Lgl1 phosphomutants as well as Lgl1-WT, and tested for the presence of α- and β-catenin in the immunoprecipitates. Neon-Lgl1-WT and Neon-Lgl1-S6A co-immunoprecipitated with α- and β-catenin. By contrast, Neon-Lgl1-S6D showed a significant reduction in its ability to form a complex with α- and β-catenin (Figure 4E). Quantification of the amount of α- and β-catenin immunoprecipitated by Neon-Lgl1-S6A or Neon-Lgl1-S6D further supports our hypothesis (Figure 4F). These results indicate that Lgl1 phosphorylation by aPKCζ regulates the ability of Lgl1 to form a complex with α- and β-catenin.

Together these results indicate that Lgl1 phosphorylation regulates not only its localization in epithelial cells, but also the interaction between Lgl1 and AJ proteins, affecting the AJs localization of E-cadherin and of α- and β-catenin, underscoring the significance of Lgl1 localization in maintaining the integrity of AJs.

### Lgl1 and Scrib affect the activation state of NMII-A

The E-cadherin-catenin complex is a key element of epithelial cell-cell adhesions that provides strong mechanical coupling between neighboring cells through association with the acto-myosin cytoskeleton ^50^. NMII regulates cell-cell adhesion by concentrating E-cadherin at contacts and activating α-catenin ^25-27^. Furthermore, NMII-A regulates β-catenin-mediated EMT ^51^ .We have previously shown that Lgl1 regulates NMII-A cellular localization in migrating cells by regulating the state of the NMII-A filament assembly ^14, 29, 30^. Since depletion of either Lgl1 or Scrib led to dissociation of E-cadherin-catenin from AJs, we tested whether Lgl1 and/or Scrib are important for the AJ localization of NMII-A. To this end, we examined the AJ localization of the NMII-A in Lgl1^KD^ and Scrib^KD^ cell lines. We found that in control cells, NMII-A appeared diffused at AJs, and in cells depletion of either Lgl1 or Scrib led to the appearance of large amounts of NMII-A in stress fibers, implying NMII-A overassembly at AJs (Figure 5A-B). To further analyze the effect of Lgl1 and Scrib on NMII-A, we acquired Z-stacks images of E-cadherin and NMII-A at AJs. Upon Lgl1 or Scrib depletion, there was a strong staining of NMII-A at AJs, further indicating the overassembly of NMII-A at AJs (Figure 5C-D). Re-expression of either Lgl1 or Scrib but not of Neon-Lgl1-S6D restored the appearance of NMII-A, as in the control cells (Figure 5C-D). These results indicate that Lgl1 and Scrib regulate NMII-A filament assembly in epithelial cells, and that in their absence, there is an increase in NMII-A associated with stress fibers. Furthermore, Lgl1 phosphorylation by aPKCζ inhibits the interaction of Lgl1 with NMII-A leading to NMII-A filament overassembly. These results are consistent with our previous findings in fibroblasts and MDA-MB-231 cells, according to which Lgl1 has an inhibitory effect on the NMII-A filament assembly and upon Lgl1 phosphorylation by aPKCζ this inhibitory effect is removed ^14, 29, 30^. Next, we investigated the complex formation of NMII-A with the E-cadherin-catenin complex. We found that endogenous NMII-A co-immunoprecipitated with endogenous α- and β-catenin (Figure 5E). This result indicates that in epithelial cells NMII-A resides in a complex with the cadherin-catenin protein complex. Repeated attempts at detecting direct interactions between NMII-A with either α- or β-catenin failed, therefore, we can state with a high degree of confidence that these proteins do not interact directly. We have previously shown that in migrating cells, Lgl1 interacts directly with NMII-A to regulate its cellular localization ^30^, leading us to postulate that Lgl1 links NMII-A to cadherin-catenin protein complex. To test this hypothesis, we immunopercipitated β- catenin from cells depleted of Lgl1 and tested for the presence of NMII-A in the immunoprecipitates. As shown in Figure 5F, NMII-A, E-cadherin, α- and β-catenin form a complex in cells expressing Lgl1. In contrast, in the absence of Lgl1, NMII-A did not form a complex with cadherin-catenin proteins. Lgl1 depletion did not affect β-catenin interaction with α-catenin or E-cadherin. These results indicate that Lgl1 serves as a linker between NMII-A and cadherin-catenin protein complex (Figure 5G).

**Figure 5:**
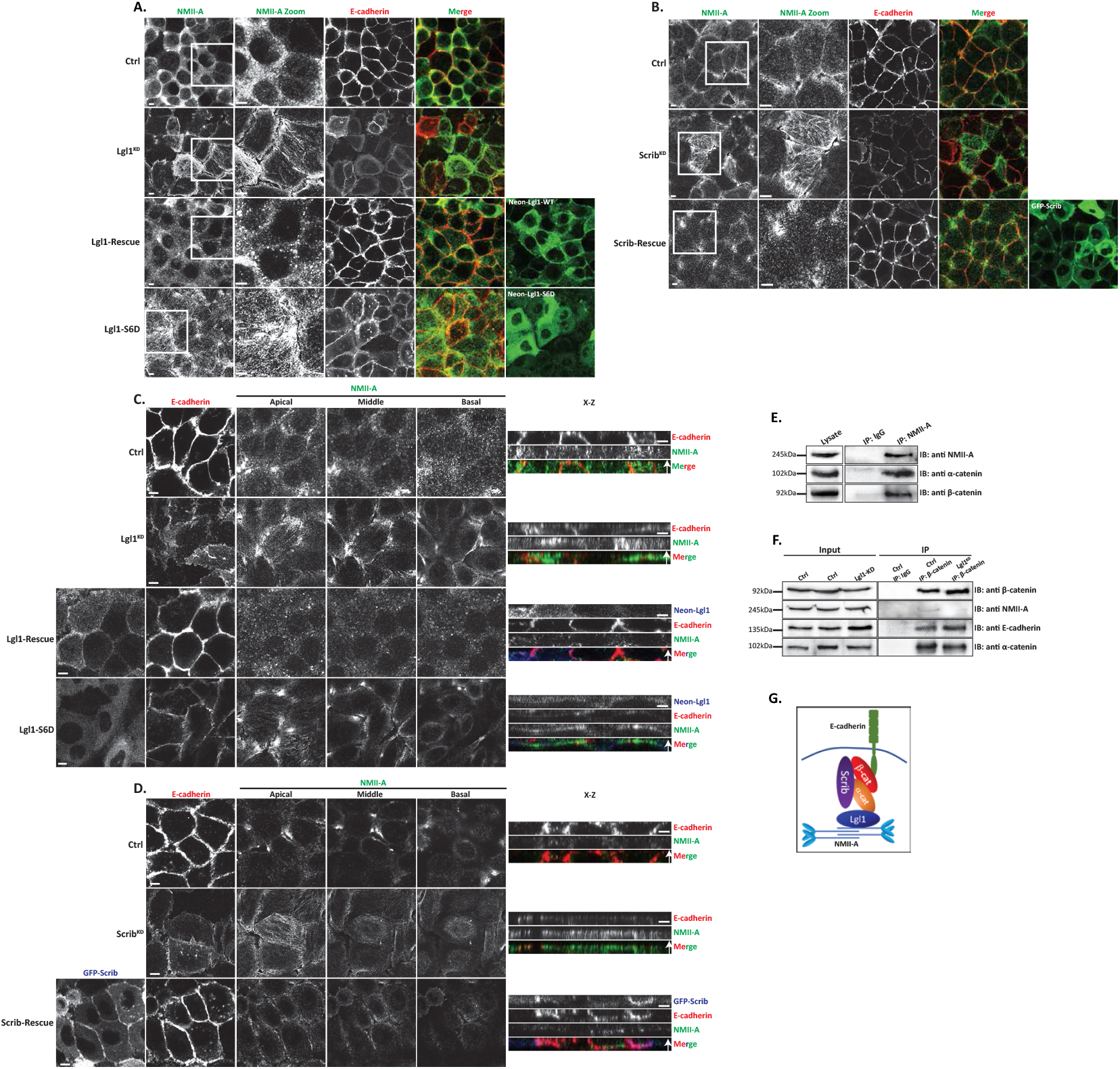
Lgl1 and Scrib regulate NMII-A at AJs. **(A)** A549 tet-shLgl1 without (Ctrl) and with Dox (Lgl1^KD^), A549 tet-shLgl1-Neon-Lgl1 with Dox (Lgl1-Rescue), and A549 tet-shLgl1-Neon-Lgl1-S6D with Dox (Lgl1-S6D) were fixed and immunostained for NMII-A and E-cadherin. **(B)** A549 tet-shscrib without (Ctrl) and with Dox (Scrib^KD^) and A549 tet-shScrib-GFP-Scrib with Dox (Scrib-Rescue) cell lines were fixed and immunostained for NMII-A and E-cadherin. Scale bar, 10 µm. White boxes indicate zoom-in on NMII-A staining. **(C)** and **(D)** Left, representative images from Z-stack of E-cadherin from the apical side and of NMII-A from apical, middle, and basal side, scale bar, 10 µm. Right, Z-stack constructed from serial optical sections along apico-basal axis. White arrow indicates apical direction along apico-basal axis in Z-stack. Scale bar, 5 µm. **(E)** A549 cell extracts were subjected to immunoprecipitation assay using NMII-A antibodies. The immunoprecipitated proteins were analyzed by IB with antibodies against NMII-A, α- and β-catenin. IgG antibody was used as a negative control. **(F)** A549 tet-shLgl1 shLgl1 without (Ctrl) and with Dox (Lgl1^KD^) cell extracts were subjected to immunoprecipitation assay using β-catenin antibodies. The immunoprecipitated proteins were analyzed by IB with antibodies against NMII-A, E-cadherin, α- and β-catenin. IgG antibody was used as a negative control. (G) A model depicting the protein complex formed by Lgl1, Scrib, E-cadherin, α- and β-catenin, and NMII-A at AJs.

The appearance of NMII-A overassembly in AJs of Lgl1^KD^ and Scrib^KD^ cells may indicate that in the absence of Lgl1 or Scrib, NMII-A is activated. To test this, we immunostained Lgl1^KD^ and Scrib^KD^ cells for Thr18/Ser19-phosphorylated MRLC (pMRLC) and acquired Z-stack images. We found that upon Lgl1 depletion, there is an increase in pMRLC in AJs (Figure 6A). By contrast, depletion of Scrib had no apparent effect on pMRLC localization at AJs (Figure 6B). Thus, Lgl1 may affect NMII activation by inhibiting MRLC phosphorylation. To further test the effect of Lgl1 and Scrib on MRLC phosphorylation, we quantified the total pMRLC in cells depleted for either Lgl1 or Scrib. Depletion of Lgl1 led to a more than two-fold increase in pMRLC (Figure 6C-D). Thus, NMII-A activation occurred in correlation with an increase of pMRLC at cell-cell junctions, further indicating that Lgl1 inhibits MRLC phosphorylation. In contrast to Lgl1, depletion of Scrib resulted in a two-fold decrease in pMRLC (Figure 6C-D), indicating that Scrib positively regulates MRLC phosphorylation. Note, staining of pMRLC does not differentiate between NMII isoforms.

**Figure 6:**
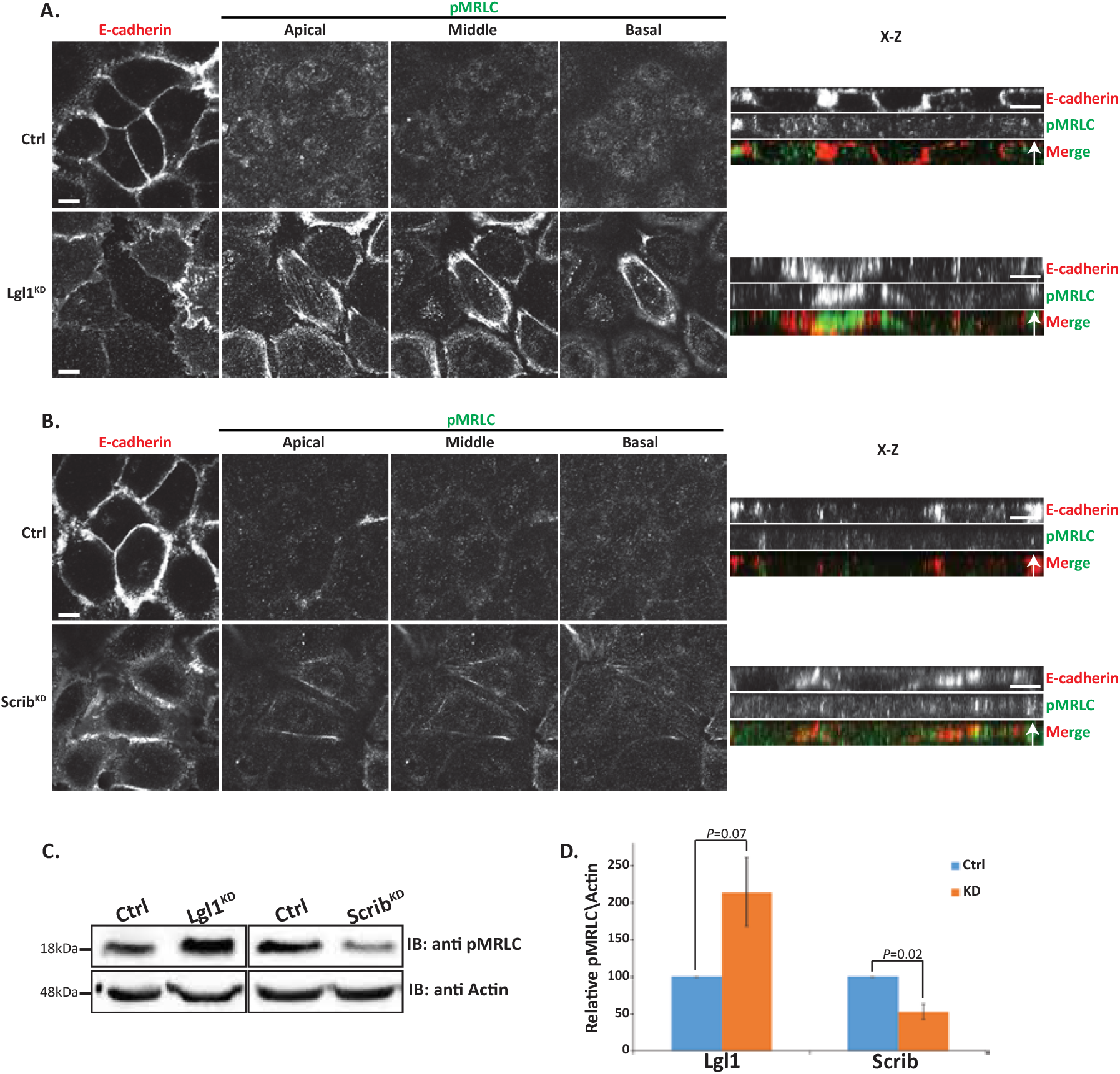
Lgl1 and Scrib regulate NMII activation at AJs. A549 tet-shLgl1 without (Ctrl) and with Dox (Lgl1^KD^), A549 tet-shscrib without (Ctrl) and with Dox (Scrib^KD^) cell lines indicated were stimulated with TGFβ for 30 min. **(A)** and **(B)** cells were fixed and immunostained for pMRLC and E-cadherin. Left, representative images from Z-stack of E-cadherin from apical side and pMRLC from apical, middle, and basal side. Scale bar, 10 µm. Right, Z-stack obtained as described above. Scale bar, 5 µm. **(C)** Levels of pMRLC in Lgl1- and Scrib-depleted cells relative to Actin. Cell lysates were subjected to IB with antibodies against pMRLC and Actin. **(D)** Quantification of amounts of pMRLC relative to Actin. Values are the mean ± SD from three independent experiments subjected to two-tailed, two-sample, and unequal-variance Student’s *t* test. Molecular weights of the proteins are indicated.

Together, these results suggest that NMII-A in epithelial A549 cells is subjected to several discrete regulation mechanisms. Lgl1 inhibits the NMII-A filament assembly and MRLC phosphorylation, whereas Scrib activates MRLC phosphorylation. In addition, because in the absence of Scrib the localization of Lgl1 to AJs is disrupted, it is possible that Scrib affects NMII-A filament assembly through its effect on Lgl1.

### TGFβ downregulates Lgl1 and Scrib expression and cellular localization

The data presented here, together with data from other labs, indicate that the localization of Lgl1 and Scrib at AJs plays an important role in AJ stabilization. Scrib forms a complex with β-catenin ^33-36^, and here we showed that Scrib forms a complex with E-cadherin and with α- and β-catenin through a direct interaction with α- and β-catenin (Figure 3). Similarly, we showed that Lgl1 forms a complex with E-cadherin and α- and β-catenin through a direct interaction of Lgl1 with α-catenin (Figure 3). Thus, Scrib and Lgl1 play important roles in AJ stabilization. AJ stability is an important factor in the EMT process because EMT is characterized by the loss of cell-cell adhesion and apical-basal cell polarity, and by increased cell motility ^52^. Therefore, we investigated whether EMT inducer, TGFβ, affects the expression of Scrib and Lgl1. To this end, we stimulated A549 cells with TGFβ for 72 hr and examined the expression levels of Scrib and Lgl1. We found that both Scrib and Lgl1 expression were downregulated during EMT program, as verified by the repression of the epithelial marker E-cadherin and the expression of the mesenchymal marker Vimentin (Figure 7A-B). Furthermore, we found that in an engineered human mammary epithelial cell line (HMLE-Twist-ER), in which the EMT program is activated by the EMT transcription factor (EMT-TF), Twist ^53, 54^, there was downregulation of the Scrib and Lgl1 expression (Figure S5A). To our knowledge, this is the first report of downregulation of Scrib and Lgl1 expression by the EMT program. These findings led us to investigate the effect of short-term exposure of A549 cells to TGFβ on the cellular localization of Scrib and Lgl1 at cell-cell contact sites. Since the localization of Lgl1 and Scrib is essential for their function, we sought to understand whether during TGFβ-induced EMT the localization of Scrib and Lgl1 is disrupted before their expression is downregulated. To this end, we examined the cellular localization of Lgl1 and Scrib in cells induced with TGFβ for 16 hr, this short-term induction did not affect the expression of Lgl1 and Scrib (Figure S5B). Note that because endogenous Lgl1 staining was weak (Figure 1), we used A549 cells stably expressing Neon-Lgl1. As shown in Figure 7C-D, in control A549 cells, endogenous Scrib, Neon-Lgl1 and E-cadherin co-localized at AJs, whereas exposure to TGFβ disrupted the localization of these proteins to AJs (Figure 7C-D), in a fashion similar to the effect of Lgl1 depletion on Scrib and E-cadherin, and *vice versa* (Figure 1). These observations were confirmed by quantification of the fluorescence intensity of Neon-Lgl1, E-cadherin and Scrib across the cell-cell junctions in A549 cells exposed to TGFβ (Figure 7E-F). Thus, TGFβ may have short- and long-term effects on AJ integrity. The short-term effect of TGFβ disrupts Lgl1 and Scrib localization at AJs, leading to disruption of E-cadherin integrity and AJ architecture; the long-term effect of TGFβ downregulates Lgl1, Scrib, and E-cadherin expression, which leads to dissolution of AJs and to EMT.

**Figure 7:**
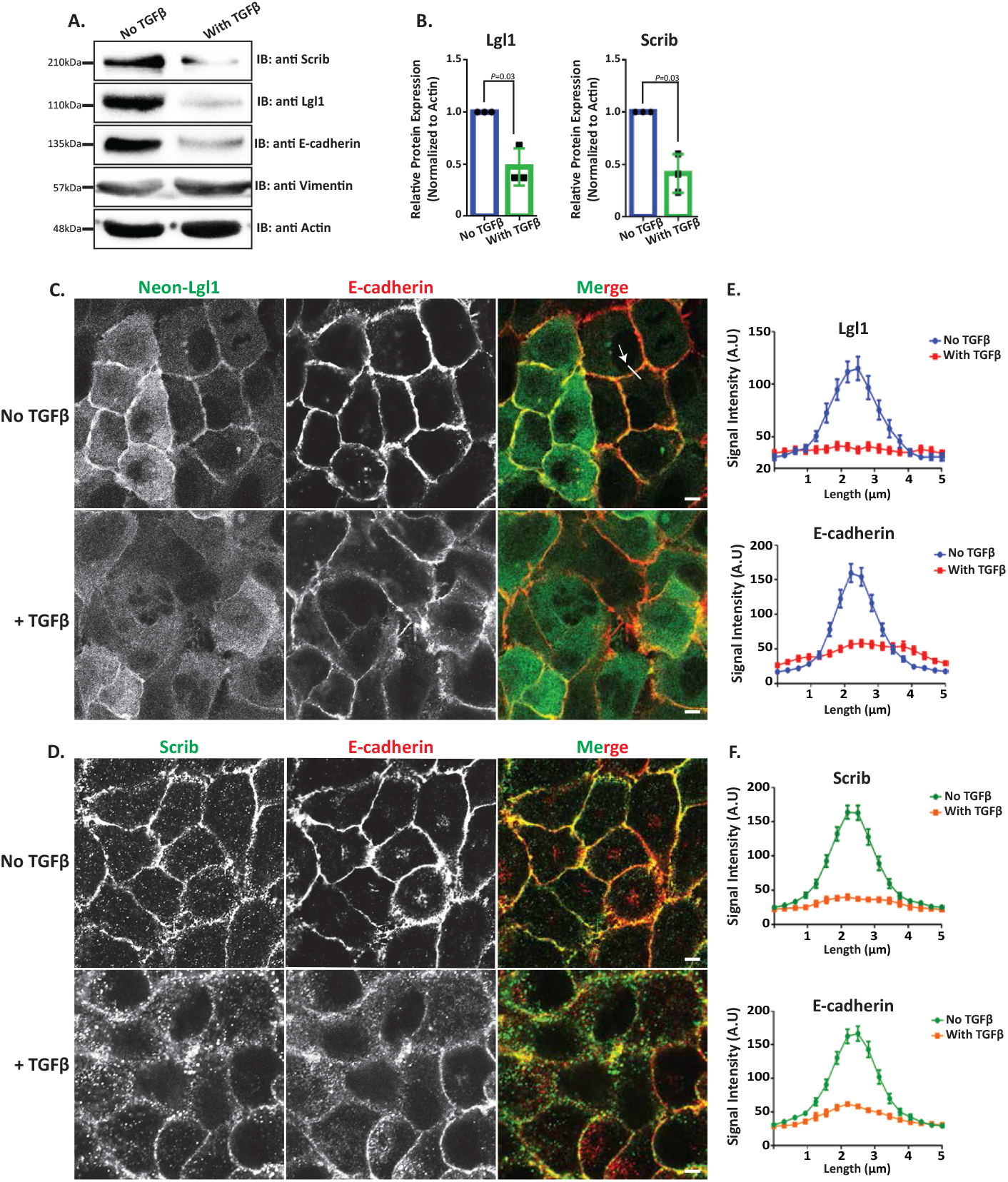
Lgl1 and Scrib are downregulated during TGFβ-induced EMT. **(A)** A549 cells were stimulated with TGFβ for 72 hr, and cell lysates were analyzed by IB with antibodies against Scrib, E-cadherin, Lgl1, Vimentin, and Actin. A549 cells without TGFβ were used as control. Molecular weights of the proteins are indicated. **(B)** Quantification of Scrib and Lgl1 protein levels with and without TGFβ relative to Actin. Values are the mean ± SD from three independent experiments subjected to two-tailed, two-sample, and unequal-variance Student’s *t* test. **(C)** and **(D)** A549 cells and A549-tet-shLgl1 (with Dox) cells expressing Neon-Lgl1 were stimulated with TGFβ for 16 h, fixed and immunostained for Scrib and E-cadherin. Scale bar, 10 µm. **(E)** and **(F)** Fluorescence intensity of endogenous E-cadherin, Scrib, and Neon-Lgl1 was measured at AJs. Results are mean ± SD, n= 30. A.U.: arbitrary units.

### Lgl1 and Scrib repress TGFβ-driven EMT

Our results to this point strongly indicate that Lgl1 and Scrib play important roles in AJ integrity by binding to the E-cadherin-catenin complex, and that TGFβ leads to the dissolution of not only E-cadherin but also of Lgl1 and Scrib from AJs. Therefore, we postulated that re-expression of Lgl1 or Scrib in cells treated with TGFβ may repress or impede the progression of TGFβ-induced EMT. To test this hypothesis, we used A549 cells that stably express GFP-Lgl1 or GFP-Scrib upon doxycycline (Dox) induction. We treated these cells with TGFβ to downregulate the endogenous Lgl1 and Scrib, then added Dox to them to induce the expression of GFP-Lgl1 or GFP-Scrib (Figure 8A). To evaluate the effect of re-expression of Lgl1 or Scrib on EMT, we determined the amounts of E-cadherin after the induction of Lgl1 or Scrib expression. Indeed, cells treated with only TGFβ presented a decrease in E-cadherin expression levels, but re-expression of Scrib or Lgl1 led to upregulation of E-cadherin (Figure 8B). Next, we quantified the E-cadherin protein levels with and without TGFβ relative to E-cadherin expression levels in control cells, normalized according to Actin (Figure 8C). Re-expression of Lgl1 led to an approximately 2.2-fold increase in E-cadherin expression in comparison to cells treated only with TGFβ. Similarly, re-expression of Scrib led to an approximately 1.5-fold increase in E-cadherin expression.

**Figure 8:**
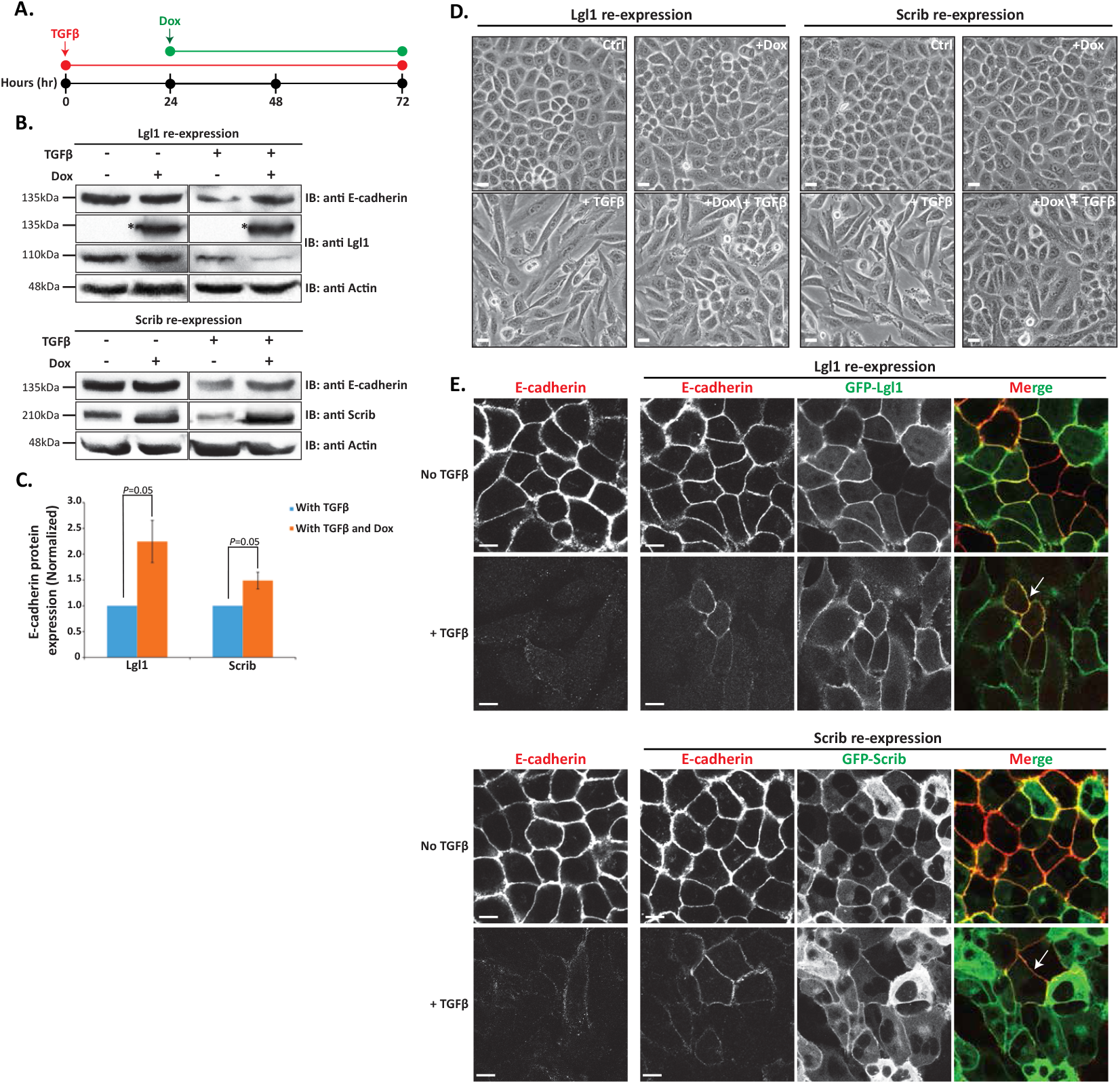
Lgl1 and Scrib re-expression impedes TGFβ-induced EMT. **(A)** Schematic representation of time-line of cell treatment with TGFβ and Dox. A549 cells expressing Dox-inducible GFP-Lgl1 or Dox-inducible GFP-Scrib were stimulated with TGFβ for 24hr followed by incubation with or without Dox for additional 48hr. **(B)** Cell lysates were analyzed by IB with antibodies against E-cadherin, Scrib or Lgl1 and Actin. Molecular weights of the proteins are indicated. *GFP-Lgl1. **(C)** Quantification of Scrib and Lgl1 protein levels with TGFβ, with or without Dox relative to control levels normalized to actin. Values are the mean ± SD from three independent experiments subjected to two-tailed, two-sample, and unequal-variance Student’s *t* test. **(D)** Phase-contrast images of the cells were taken by fluorescence microscopy. Scale bar, 20 µm. **(E)** Cells treated as in A were fixed and immunostained for E-cadherin. E-cadherin, GFP-Lgl1 and GFP-Scrib were visualized by confocal microscope. Scale bar, 10 µm. Arrows indicate co-localization of E-cadherin and GFP-Lgl1 or GFP-Scrib at AJs.

To further study the effect of Lgl1 and Scrib re-expression on the EMT process, we analyzed the morphology of A549 monolayers treated with TGFβ. Untreated cells had the morphology of an epithelial cell sheet, with apparent AJs (Figure 8D). Upon TGFβ treatment, these cells displayed a polarized spindle-shaped morphology similar to that of cells undergoing TGFβ-mediated EMT (Figure 8D). However, upon induction of Lgl1 or Scrib expression after EMT initiation, these cells became clustered and displayed a more pronounced epithelioid morphology, similar to that of cells without TGFβ indicating obstruction in EMT program (Figure 8D). To further understand the effect of induction of Lgl1 or Scrib on EMT, we immunostained these cells for E-cadherin. Induction of Lgl1 or Scrib expression after EMT initiation partially restored the localization of E-cadherin at AJs, as well as the localization of GFP-Lgl1 or GFP-Scrib (Figure 8E). Cytoplasmic localization of GFP-Lgl1 or GFP-Scrib may be attributed to their overexpression. Together, these results provide evidence that Scrib and Lg1 re-expression upon TGFβ-mediated EMT induction may impede progression.

## Discussion

In this study, we explored the role of Lgl1 and Scrib in maintaining AJ integrity. Our results show that both Scrib and Lgl1 are important for AJ integrity, and that the localization of Scrib and Lgl1 at AJs is interdependent, as we previously observed in migrating cells ^14^. Thus, Scrib and Lgl1 may perform their function at AJs as a complex, similarly to their function in cell migration ^14^. Additionally, through an array of biochemical assays we were able to show that the Scrib PDZ domains form a complex with the E-cadherin-catenin complex protein through a direct interaction with α- and β-catenin. Several studies have shown that Scrib forms a complex with β-catenin ^33-36^, but none of these studies demonstrated a direct interaction between Scrib and β-catenin, and provided no indication of Scrib-α-catenin interaction. Therefore, our findings provide new insights into the mechanism of AJ architecture. Scrib is a large scaffold protein that consists of four PDZ domains, each with distinct protein interactions ^12^. Therefore, it is conceivable that at AJs, Scrib interacts simultaneously with both α- and β-catenin. In mouse brain cells, Lgl1 has been reported to be essential in maintaining AJs ^38^, but the role of Lgl1 in cell-cell contact in epithelial cells is poorly understood. We show that similarly to Scrib, Lgl1 forms a complex with the E-cadherin-catenin complex protein, but Lgl1 interacts directly only with α-catenin. Thus, Lgl1 and Scrib may affect the E-cadherin-catenin complex to regulate AJ integrity in a different manner, and may be considered as part of the E-cadherin-catenin complex proteins. A similar conclusion has been reached for Scrib, which was shown to be a regulator of the E-cadherin-catenin complex in MDCK cells ^32, 45^. Based on our results, it is possible to conclude that E-cadherin also regulates Scrib and Lgl1 AJ localization, as TGFβ-induced downregulation of E-cadherin, prior to downregulation of Scrib and Lgl1 expression, led to disruption of Scrib and Lgl1 localization. These results are consistent with the findings that depletion of E-cadherin phenocopies the effects of Scrib depletion in MDCK cells at cell-cell contact sites ^32^, demonstrating an interdependent relationship between Scrib and E-cadherin. We propose that Scrib and Lgl1 form an important part of AJs and serve to stabilize the E-cadherin-catenin complex required to maintain AJ integrity (Figure 3H).

aPKCζ phosphorylates and inactivates Lgl1 at the apical side, regulating Lgl1 cellular localization and protein-protein interactions ^30, 48, 55, 56^. We found that Lgl1 phosphorylation by aPKCζ inhibits its interaction with α- and β-catenin, and that phosphomimetic Lgl1 was unable to rescue the AJ phenotype observed in Lgl1-depleted cells. These are consistent with the findings that aPKCζ phosphorylation of Lgl1 negatively regulated Lgl1 interaction with N-cadherin ^38^. Together, these findings reveal the significance of Lgl1 membrane association in regulating AJ architecture and add functional relevance to aPKCζ phosphorylation of Lgl1. We speculate that aPKCζ regulates Lgl1 AJ localization during cell-cell contact formation, and that during EMT, aPKCζ is activated leading to Lgl1 phosphorylation and dissociation of Lgl1 from AJs, resulting in E-cadherin-catenin dissociation form AJs and to AJ dissolution (Figure 9A). In addition, because aPKCζ is part of the Par polarity complex ^57^, our results provide a link between the Scribble and the Par polarity complexes at AJs ^58^. Therefore, we hypothesize that these polarity complex proteins operate jointly to maintain AJs, in addition to their function in maintaining apico-basal cell polarity.

**Figure 9:**
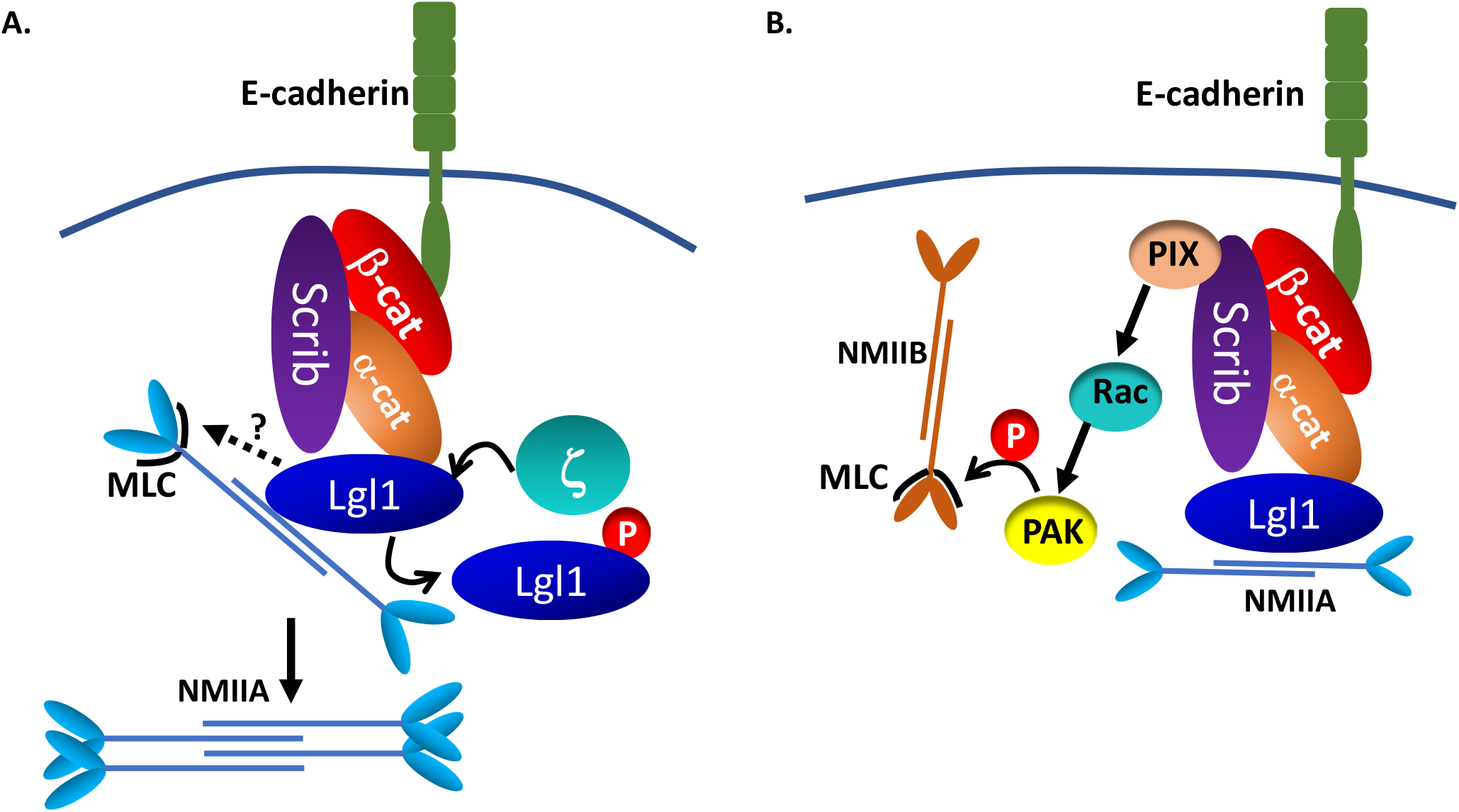
A model depicting the mechanism by which Lgl1 and Scrib maintain AJ integrity. **(A)** Scrib and Lgl1 stabilize AJs by forming a complex with E-cadherin-catenin by direct interaction with α- and β-catenin, as well as with NMII-A. Lgl1 regulates NMII-A filament assembly by binding to NMII-A rod, thus regulating the amounts of NMII-A at AJs. aPKCζ phosphorylation of Lgl1 leads to Lgl1 removal from α- and β-catenin and removal from NMII-A leading to NMII-A filament assembly. Lgl1 also affects MRLC phosphorylation by unknown mechanism. **(B)** Scrib affects MRLC phosphorylation through binding to NMII-B and/or through the Scrib → β-Pix → Rac → PAK → pMRLC pathway.

It has been previously shown that NMII-A is important for AJ assembly and preservation ^27, 51^. Our findings indicate that NMII-A resided in a complex with E-cadherin and with α- and β-catenin, but we did not detect a direct interaction between these proteins. Because NMII-A interacts directly with Lgl1 ^30^, we propose that NMII-A forms a complex with E-cadherin-catenin through its direct interaction with Lgl1 (Figure 9A). Previously, we have shown that Lgl1 regulates NMII-A filament assembly in migrating cells ^30^. It is likely that epithelial cells are regulated by a similar mechanism (Figure 9A). Indeed, we found that in the absence of Lgl1, there are clusters of NMII-A that may point at NMII-A filament over-assembly. Furthermore, NMII-A activation upon Lgl1 depletion was also suggested by the increased levels of pMRLC. Notably, in a recent study, elevated pMRLC levels induced by RhoA activation were reported in α-catenin knock-out cells, ^59^. We propose that Lgl1 binding to the rod domain of NMII-A in epithelial cells directly regulates NMII-A filament assembly, and that MRLC phosphorylation is regulated indirectly by Lgl1 (Figure 9A). We further found that depletion of Scrib also led to NMII-A filament over-assembly, although Scrib does not bind NMII-A ^14^. We propose that Scrib does not affect NMII-A directly, but rather through Lgl1, as Scrib depletion results in Lgl1 mis-localization at AJs. Thus, it is plausible that Scrib depletion disrupts Lgl1 localization to AJs, relieving the Lgl1 inhibition of NMII-A, and leading to over assembly of NMII-A filaments (Figure 9A). This hypothesis is supported by our finding that phosphomimetic Lgl1, which fails to rescue the Lgl1 depletion phenotype and does not bind NMII-A ^29^, also exhibited NMII-A over-assembly, leading to the conclusion that Lgl1 localization at AJs is pivotal for NMII-A regulation. Note that expression of E-cadherin-α-catenin, a chimeric that lacks β-catenin binding sites but can connect to the actin cytoskeleton, was able to restore AJ integrity in cells lacking NMII-A or Scrib in a similar manner ^27, 32^, linking Scrib and NMII-A to the same regulating complex of AJs, possibly through Lgl1. In contrast to Lgl1-depleted cells, in Scrib-depleted cells we observed a decrease in MRLC phosphorylation. MRLC are not NMII isoform specific, therefore, MRLC may bind to all NMII isoforms to regulate their activity ^60^, therefore, an increase in MRLC phosphorylation may lead to the activation of any NMII isoform. Furthermore, we have previously shown that Scrib forms a complex with NMII-B specifically ^14^. In addition, NMII isoforms play distinct roles at AJs, NMII-A plays a role in E-cadherin clustering, while NMII-B provides stability for the actin belt ^61^. Thus, we propose that by binding to NMII-B, Scrib may promote phosphorylation of MRLC that are associated with NMII-B, and upon Scrib depletion, there is a decrease in MRLC phosphorylation, leading to NMII-B inactivation affecting the actomyosin belt at AJs (Figure 9B). Scrib may affect MRLC phosphorylation through its regulatory effect on the kinase PAK, which presumably phosphorylates MRLC (Figure 9B) ^62^. It has been shown that Scrib directly binds the GEF, β-Pix, which activates Rac ^13^. The absence of Scrib causes a significant decrease in Rac activity ^63^, leading to PAK inactivation. Thus, we hypothesize that Scrib depletion causes a decrease in Rac activity and inactivation of PAK, leading to a reduction in MRLC phosphorylation and inhibition of NMII-B. In this way, Scrib and Lgl1 differentially regulate NMII-A and NMII-B at AJs.

Downregulation and mis-localization of Scrib and Lgl1, among other polarity proteins, has been associated with cancer progression in carcinomas ^64^. Additionally, Scrib and Lgl1 were found to function as tumor suppressors ^65, 66^. Here we show that TGFβ−induced EMT led to a significant downregulation of Scrib and Lgl1 protein expression, indicating that in addition to E-cadherin, Scrib and Lgl1 are also targeted during the EMT program. We found that TGFβ-induced EMT has short- and long-term effects on Scrib and Lgl1 AJs localization. Prior to TGFβ-induced Scrib and Lgl1 downregulation, TGFβ leads to Scrib and Lgl1 mis-localization. We propose that the downregulation and/or mis-localization of Scrib and Lgl1 observed in many types of cancer contribute to EMT by affecting E-cadherin-catenin complex stability, leading to AJ dissociation. We cannot rule out that at least in part, the short-term effect of TGFβ on Scrib and Lgl1 is the result of E-cadherin downregulation. In an attempt to understand the mechanism that leads to Scrib and Lgl1 downregulation by TGFβ induced EMT, we treated A549 cells with the proteasome inhibitor MG132, with and without TGFβ, and observed Scrib and Lgl1 protein levels. The MG132 inhibitor did not affect Scrib or Lgl1 protein downregulation by TGFβ (data not shown). Indicating that downregulation of these proteins is not a result of protein degradation or instability induced by the EMT program, and suggesting regulation at mRNA levels. TGFβ induces EMT by activation of three families of EMT-TFs: the Snail, ZEB, and bHLH families ^67^. It has been previously reported that Lgl2 is a target for ZEB1 and Snail, as depletion of either EMT-TFs caused upregulation of Lgl2 expression ^68-70^. It has also been shown that Snail expression leads to Scrib mis-localization, without affecting protein expression ^71, 72^. Similarly, Snail has been reported to disrupt localization of both the Crumbs and Par complexes at tight junctions, and to repress expression of only Crumbs ^73^, indicating uncoupling between localization and expression regulation by EMT-TFs. Presently, the EMT-TFs that may regulate Scrib and Lgl1 expression in A549 cells are unknown.

Our results support the role of Scrib as a tumor suppressor. At the same time, Scrib has also been reported to be over-expressed in cancer cells and behave as an oncogenic protein ^74, 75^. These findings suggest that Scrib plays a context-specific role in epithelial tissue, both as a tumor suppressor and as an oncogene ^76^. Furthermore, our results do not exclude the possibility that Scrib re-expression occurs during metastasis, as EMT is a transient event, and that the converse mesenchymal-to-epithelial transition also occurs in the course of metastasis ^77^.

Re-expression of Scrib and Lgl1 during EMT was able to impede EMT progression by partially restoring E-cadherin expression and localization. Similarly, it was reported that re-expression of exogenous Crumbs3 in Snail-induced EMT partially restored cell-cell junctions ^73^. Furthermore, increased expression of Scrib has been shown to promote tight junction formation through downregulation of the EMT inducer, ZEB ^78^. Loss of cell polarity is considered to be mainly the passive result of EMT inducing signals that enable loss of epithelial characteristics. The results presented in this study raise the possibility that the Scribble polarity complex actively inhibits EMT, possibly by regulating, for example, the nuclear localization of EMT-TFs, as was shown for Lgl2. Lgl2 reduces the nuclear localization of Snail, binding Snail to its target promotor and inhibiting EMT. It is plausible that Scrib and Lgl1 inhibit EMT-TFs function, thus inhibiting EMT.

In conclusion, our findings identify Scrib and Lgl1 as essential for AJ integrity and cytoskeleton arrangement in epithelial cells, making them an obvious target for downregulation during EMT induction. Our findings, together with those of our previous study ^14^, reveal the common and individual contribution of Scrib and Lgl1 to cell adhesion and migration, providing a foundation for understanding their disruption during metastasis.

## Data Availability Statement (DAS)

The data and materials that support the results or analyses presented in this paper are freely available upon request.

## Acknowledgment

We thank the Pawson’s lab (Mount Sinai Hospital, Toronto, Canada) for the Lgl1 construct, Patrick Humbert and Helena Richardson (La Trobe University, Melbourne, Australia) for Scrib constructs, Robert S. Adelstein (NIH) for NMII constructs, Yinon Ben-Neria (Hebrew University) for β-catenin constructs, Cara Gottardi (Northwestern University) for GST-α-catenin, and Yael Feinstein Rotkopf for technical assistance with the microscopy work. This study was supported by the Israel Science Foundation (Grant No. 745/17), Israel Cancer Research Foundation, and Israel Cancer Association (Grant No. 20140082). S.R. holds the Daniel G. Miller Chair in Cancer Research.

