## Supplemental Figures for "Scribble, Lgl1, and myosin IIA interact with α/β-catenin to maintain epithelial junction integrity"

**Supplemental Figure Legends**

**Supplemental Figure S1:** The expression of Scrib, Lgl1, E-cadherin, and α- and β-catenin were analyzed in Lgl1- **(A)** and Scrib- **(B)** depleted cell lines, as well as in cells expressing Neon-Lgl1 proteins **(C)** or GFP-Scrib **(D)**. Actin served as a loading control. Black and green arrows in C, indicate endogenous and Neon fusion proteins, respectively. Molecular weights of the proteins are indicated.

**Supplemental Figure S2:** Dot-plot of signal intensity of junctional protein in comparison to cytoplasmic protein of E-cadherin **(A),** Scrib **(B),** α-catenin **(C),** and β-catenin **(D)** in the indicated Lgl1 cell lines. Values are the mean ± SD from three independent experiments subjected to ANOVA, with a *post hoc* test. *ns*: not significant.

**Supplemental Figure S3:** Dot-plot of signal intensity of junctional protein in comparison to cytoplasmic protein of E-cadherin **(A),** Lgl1 **(B),** α-catenin **(C),** and β-catenin **(D)** in the indicated Scrib cell lines. Values are the mean ± SD from three independent experiments subjected to ANOVA, with a post *hoc test*. *ns*: not significant.

**Supplemental Figure S4: (A)** A549 cell extracts were subjected to co-IP assay using Scrib antibody. The immunoprecipitated proteins were analyzed by IB with antibodies against Scrib and E-cadherin. IgG was used as negative control. **(B)** MBP-Lgl1 and GST only or GST-β-catenin were subjected to PD assay. MBP-Lgl1 was analyzed by IB with antibody against MBP-tag, and GST proteins were analyzed by Ponceau S staining. Molecular weights of the proteins are indicated.

**Supplemental Figure S5: (A)** HMLE-Twist-ER cells were induced by 4-hydroxytamoxifen (OHT) for the indicated time points, and cell lysates were analyzed by IB with antibodies against Scrib and Lgl1. Actin served as a loading control. **(B)** A549 cells were incubated with TGFβ for 16 h, and cell lysates were analyzed by IB with antibodies against Scrib, E-cadherin, and Lgl1. Actin served as a loading control. Molecular weights of the proteins are indicated.

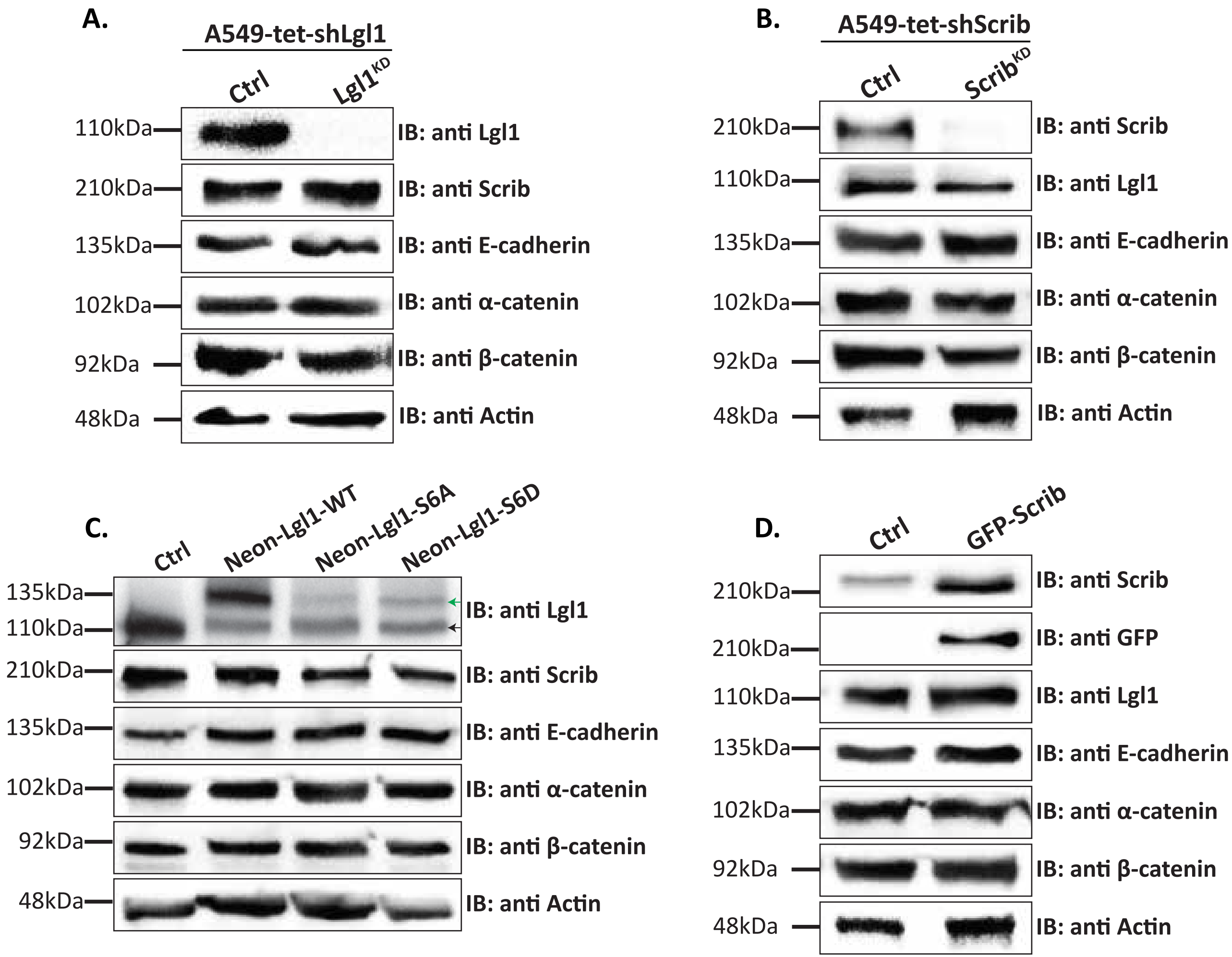


**Supplemental Figure S1:** The expression of Scrib, Lgl1, E-cadherin, and α- and β-catenin were analyzed in Lgl1- **(A)** and Scrib- **(B)** depleted cell lines, as well as in cells expressing Neon-Lgl1 proteins **(C)** or GFP-Scrib **(D)**. Actin served as a loading control. Black and green arrows in **C**, indicate endogenous and Neon fusion proteins, respectively. Molecular weights of the proteins are indicated.

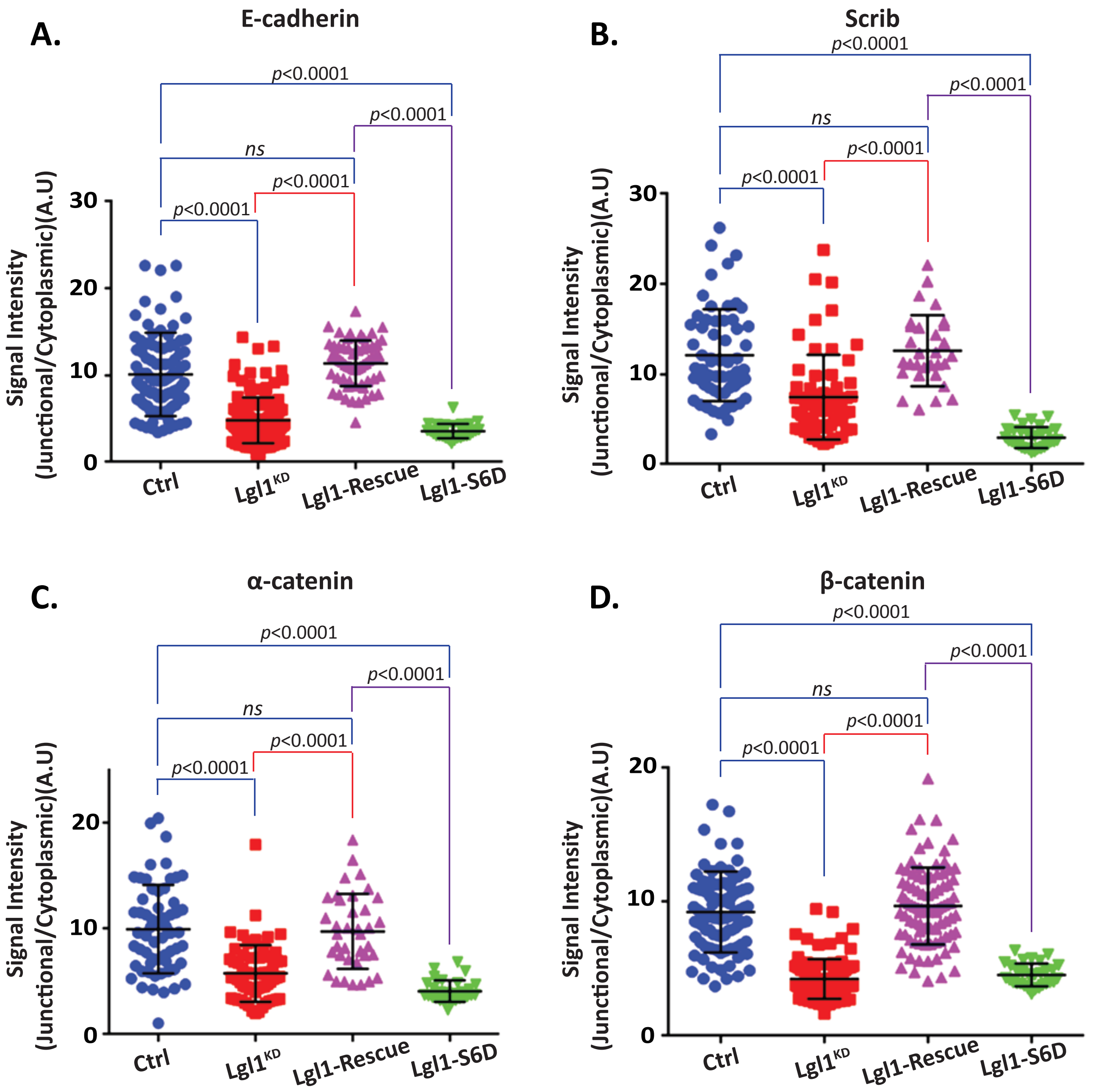


**Supplemental Figure S2:** Dot-plot of signal intensity of junctional protein in comparison to cytoplasmic protein of E-cadherin **(A)**, Scrib **(B)**, α-catenin **(C)**, and β-catenin **(D)** in the indicated Lgl1 cell lines. Values are the mean ± SD from three independent experiments subjected to ANOVA, with a post hoc test. *ns*: not significant.

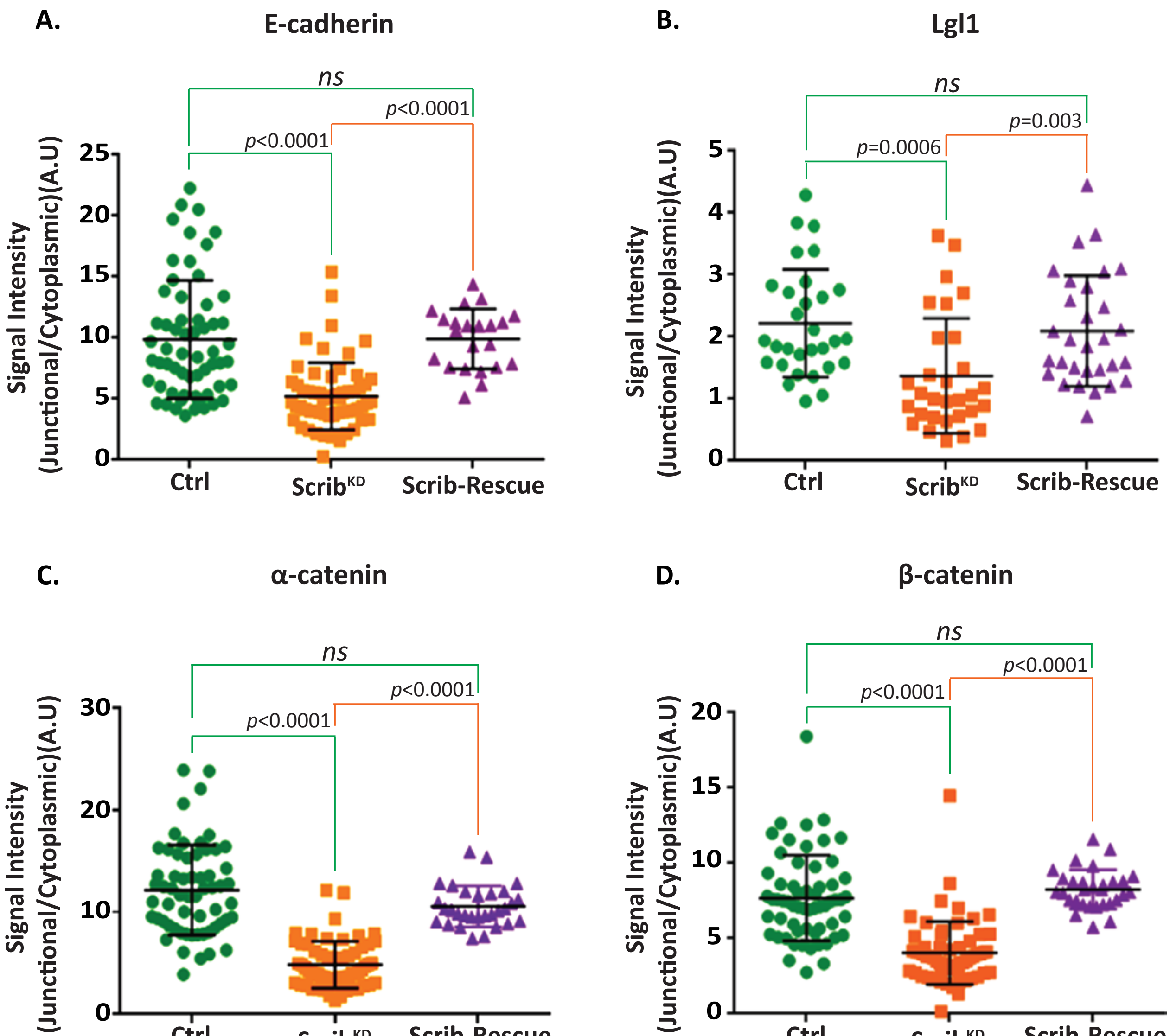


**Supplemental Figure S3:** Dot-plot of signal intensity of junctional protein in comparison to cytoplasmic protein of E-cadherin **(A)**, Lgl1 **(B)**, α-catenin **(C)**, and β-catenin **(D)** in the indicated Scrib cell lines. Values are the mean ± SD from three independent experiments subjected to ANOVA, with a post hoc test. *ns*: not significant.

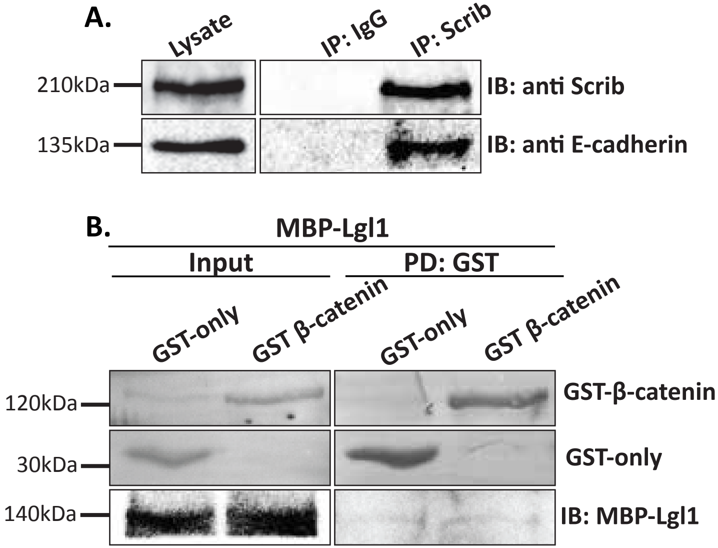


**Supplemental Figure S4: (A)** A549 cell extracts were subjected to co-IP assay using Scrib antibody. The immunoprecipitated proteins were analyzed by IB with antibodies against Scrib and E-cadherin. IgG was used as negative control. **(B)** MBP-Lgl1 and GST only or GST-β-catenin were subjected to PD assay. MBP-Lgl1 was analyzed by IB with antibody against MBP-tag, and GST proteins were analyzed by Ponceau S staining. Molecular weights of the proteins are indicated.

A.

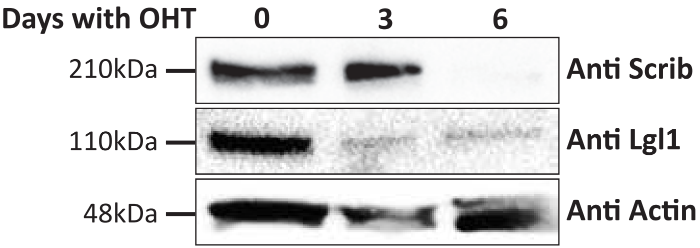


B.

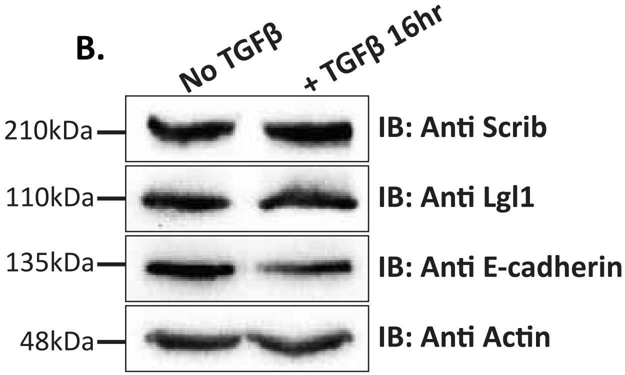


**Supplemental Figure S5: (A)** HMLE-Twist-ER cells were induced by 4-hydroxytamoxifen (OHT) for the indicated time points, and cell lysates were analyzed by IB with antibodies against Scrib and Lgl1. Actin served as a loading control. **(B)** A549 cells were incubated with TGFβ for 16 h, and cell lysates were analyzed by IB with antibodies against Scrib, E-cadherin, and Lgl1. Actin served as a loading control. Molecular weights of the proteins are
